# Zinc-finger (ZiF) fold secreted effectors form a functionally diverse family across lineages of the blast fungus *Magnaporthe oryzae*

**DOI:** 10.1101/2023.10.18.562914

**Authors:** Juan Carlos De la Concepcion, Thorsten Langner, Koki Fujisaki, Vincent Were, Xia Yan, Anson Ho Ching Lam, Indira Saado, Helen J. Brabham, Joe Win, Kentaro Yoshida, Nicholas J. Talbot, Ryohei Terauchi, Sophien Kamoun, Mark J. Banfield

**Affiliations:** Department of Biochemistry and Metabolism, John Innes Centre, Norwich Research Park, Norwich, NR4 7UH, UK; The Sainsbury Laboratory, University of East Anglia, Norwich Research Park, Norwich, NR4 7UH, UK; Gregor Mendel Institute of Molecular Plant Biology, Austrian Academy of Sciences, Vienna, 1030, Austria; Department of Molecular Biology, Max Planck Institute for Developmental Biology, Tubingen, Germany; Division of Genomics and Breeding, Iwate Biotechnology Research Center, Iwate 024-0003, Japan; Department of Cell and Developmental Biology, John Innes Centre, Norwich Research Park, Norwich, NR4 7UH UK; Laboratory of Plant Genetics, Graduate School of Agriculture, Kyoto University, Kyoto 606-8502, Japan; Laboratory of Crop Evolution, Graduate School of Agriculture, Kyoto University, Kyoto, 606-8501, Japan

## Abstract

Filamentous plant pathogens deliver effector proteins into host cells to suppress host defence responses and manipulate metabolic processes to support colonization. Understanding the evolution and molecular function of these effectors provides knowledge about pathogenesis and can suggest novel strategies to reduce damage caused by pathogens. However, effector proteins are highly variable, share weak sequence similarity and, although they can be grouped according to their structure, only a few structurally conserved effector families have been functionally characterized to date. Here, we demonstrate that Zinc-finger fold (ZiF) secreted proteins form a functionally diverse effector family in the blast fungus *Magnaporthe oryzae*. This family relies on the Zinc-finger motif for protein stability and is ubiquitously present, forming different effector tribes in blast fungus lineages infecting 13 different host species. Homologs of the canonical ZiF effector, AVR-Pii from rice infecting isolates, are present in multiple *M. oryzae* lineages, and the wheat infecting strains of the fungus, for example, possess an allele that also binds host Exo70 proteins and activates the immune receptor Pii. Furthermore, ZiF tribes vary in the host Exo70 proteins they bind, indicating functional diversification and an intricate effector/host interactome. Altogether, we uncovered a new effector family with a common protein fold that has functionally diversified in lineages of *M. oryzae*. This work expands our understanding of the diversity of *M. oryzae* effectors, the molecular basis of plant pathogenesis and may ultimately facilitate the development of new sources for pathogen resistance.

**Author Summary:** Diseases caused by filamentous plant pathogens impact global food production, leading to severe economic and humanitarian consequences. These pathogens secrete hundreds of effectors inside the host to alter cellular processes and to promote infection and disease. Effector proteins have weak or no sequence similarity but can be grouped in structural families based on conserved protein folds. However, very few conserved effector families have been functionally characterized. We have identified a family of effectors with a shared Zinc-finger protein fold (ZiF) that is present in lineages of the blast fungus *Magnaporthe oryzae* that can, collectively, infect 13 different grasses. We characterized the binding of a sub-set of these proteins to putative Exo70 host targets and showed they can be recognized by the plant immune system. Furthermore, we show that other ZiF effectors do not bind Exo70 targets, suggesting functional specialization within this effector family for alternative interactors. These findings shed light on the diversity of effectors and their molecular functions, as well as potentially leading to the development of new sources of blast disease resistance in the future.

## Introduction

Pathogens have evolved sophisticated strategies to overcome host defences and manipulate cellular pathways to promote their proliferation. One such strategy is to deploy effector proteins into the host. These effectors carry out multiple functions within the host cell, including plant immune suppression and metabolic manipulation (Sanchez-Vallet et al., 2018). In general, effectors can be very diverse, sharing low or no sequence similarity. Despite their intrinsic variability, some effectors share similar protein folds and can be grouped into structurally related families (Derbyshire and Raffaele, 2023; Franceschetti et al., 2017; Seong and Krasileva, 2023). Fungal effectors are particularly diverse, and it is challenging to classify and validate putative effectors encoded in fungal genomes (Lovelace et al., 2023; Varden et al., 2017). Elucidation of effector structures has revealed how fungal effectors with low amino acid sequence conservation can share the same protein fold (de Guillen et al., 2015). Recently, experimental information (or structural predictions) of fungal effector folds have guided genome-wide searches defining structural families (de Guillen et al., 2015; Lazar et al., 2022; Le Naour-Vernet et al., 2023; Rocafort et al., 2022; Seong and Krasileva, 2023; Teulet et al., 2023; Yu et al., 2022). However, although effector proteins can be classified according to their predicted structure, only a few structurally conserved effector families have been functionally characterized to date.

Blast disease, caused by the ascomycete fungus *Magnaporthe oryzae*, affects a wide variety of grasses including staple food crops such as rice, barley, and wheat in addition to many other grass species causing significant losses in food production required to feed the world’s population (Eseola et al., 2021; Latorre et al., 2023; Wilson and Talbot, 2009). The blast fungus is thought to be undergoing incipient speciation into genetically distinct lineages that are typically associated with a single grass host genus (Gladieux et al., 2018; Thierry et al., 2022). *M. oryzae* infects multiple wild grasses that often occur in close proximity to crops, acting as a pathogen reservoir (Barragan et al., 2022) that, together with the potential for host jumps and reassortment of the effector repertoire (Bentham et al., 2021; Inoue et al., 2017), makes this pathogen a significant threat to food security even in areas where blast disease is not endemic (Barragan et al., 2022). Therefore, understanding the virulence mechanisms and effector biology of *M. oryzae* is important to achieve resistance and develop new strategies to mitigate its economic impact (Cadiou et al., 2023; Zdrzałek et al., 2023).

The genome of *M. oryzae* encodes hundreds of putative effectors (Dean et al., 2005). The rice blast strain Guy11, for example, has at least 546 effector proteins (Yan et al., 2023). To date, most structurally characterized blast fungus effectors share the same protein fold. Structural studies of the effectors AVR-Pik, AVR1-CO39, AVR-Pia, AVR-Pizt and AVR-Pib revealed they all share a six-strand β-sandwich fold, comprised of two antiparallel β-sheets (de Guillen et al., 2015; De la Concepcion et al., 2018; Ortiz et al., 2017; Zhang et al., 2013). This was designated as the MAX (for Magnaporthe Avrs and ToxB like) effector fold and was used to define a diverse and expanded effector family in blast genomes comprising 5 to 10% of all putative effectors (de Guillen et al., 2015).

Studies focused on the biology of MAX effectors have demonstrated how these MAX effectors target host proteins (Bentham et al., 2021; Maidment et al., 2021; Oikawa et al., 2020), how they are detected by the plant immune system (De la Concepcion et al., 2018; De la Concepcion et al., 2021a; Guo et al., 2018; Maqbool et al., 2015; Ortiz et al., 2017), and how this interaction has shaped the evolution of host immunity (Bialas et al., 2021; De la Concepcion et al., 2021b). These studies have led to the first steps in the development of synthetic intracellular immune receptors with bespoke pathogen recognition (Bentham et al., 2022; Cesari et al., 2022; De la Concepcion et al., 2019; Kourelis et al., 2023; Liu et al., 2021; Maidment et al., 2023), a goal long pursued in plant biotechnology (Cadiou et al., 2023; Rodriguez-Moreno et al., 2017; Zdrzałek et al., 2023). Despite this progress, studies addressing functional characterization and experimental determination of non-MAX effectors from *M. oryzae* have remained somewhat behind in comparison to MAX effectors.

Recently, we determined the structure of the non-MAX effector AVR-Pii in complex with its host target (De la Concepcion et al., 2022). AVR-Pii is a secreted small effector protein that binds to two Exo70 proteins, OsExo70F2 and OsExo70F3, both putative components of the rice exocyst complex (De la Concepcion et al., 2022; Fujisaki et al., 2015). The physical interaction between AVR-Pii and these Exo70 proteins is recognized by the rice Pii NLR pair (Pii-1 and Pii-2), triggering immune responses that halt colonization by *M. oryzae* (Fujisaki et al., 2015; Fujisaki et al., 2017). Even though their molecular cloning was reported at the same time (Białas et al., 2018; Yoshida et al., 2009), AVR-Pii remained understudied compared with the MAX effectors AVR-Pik and AVR-Pia. This is most likely because this effector is absent in most rice blast isolates and was not found in preliminary effector searches in isolates infecting other cereals (Latorre et al., 2020; Yoshida et al., 2016).

The structure of AVR-Pii bound to OsExo70F2 identified a new effector/target molecular interface with the potential to suggest the virulence-associated mechanism of this effector and inform the development of bespoke resistance by engineering effector recognition in NLRs (De la Concepcion et al., 2022). Interestingly, the structure also revealed a novel effector fold comprising a Zinc-finger domain, coined the Zinc-finger effector fold (ZiF), which is distinct in sequence and structure from the MAX fold previously identified for Magnaporthe effectors (De la Concepcion et al., 2022). Understanding the diversity and structure / function relationships of ZiF effectors is necessary to guide plant breeding approaches as well as bioengineering of novel disease resistance genes.

Here, we tested the importance of residues forming the Zinc-finger in AVR-Pii function and folding, as a general model for ZiF effectors. We used the structural information of the AVR-Pii fold to guide a Hidden Markov Model search into the genomes of *M. oryzae* infecting diverse grasses to identify effectors sharing the ZiF fold. We report and categorize the presence of a family of ZiF effector proteins prevalent across blast lineages and show that different ZiF effector tribes likely target different host proteins, implying functional diversity. We show that, just like rice blast AVR-Pii, some wheat blast ZiF effectors bind host Exo70s and can be recognized by Pii resistance, establishing a potential novel approach for developing disease resistance against the emerging wheat blast pathogen.

## Results

### The Zinc-finger motif of AVR-Pii is required for protein stability and binding to host Exo70 proteins

The recently determined structure of AVR-Pii revealed a new fold for a *M. oryzae* effector based on a Zinc-finger formed by residues Cys51, Cys54, His67 and Cys69 (De la Concepcion et al., 2022). Although not directly involved in the effector/target binding interface, random mutagenesis assays suggest that mutating these residues has a negative effect on interaction with the host target OsExo70F3 (De la Concepcion et al., 2022). To better understand the role of the AVR-Pii Zinc-finger motif, we mutated residues Cys51, Cys54 and His67 to Alanine (AVR-Pii^CCH^) and tested for interaction with rice Exo70 targets, OsExo70F2 and OsExo70F3, by Yeast-Two-Hybrid (Y2H) and *in planta* co-immunoprecipitation (Co-IP) **(Figure 1)**.

**Figure 1.**
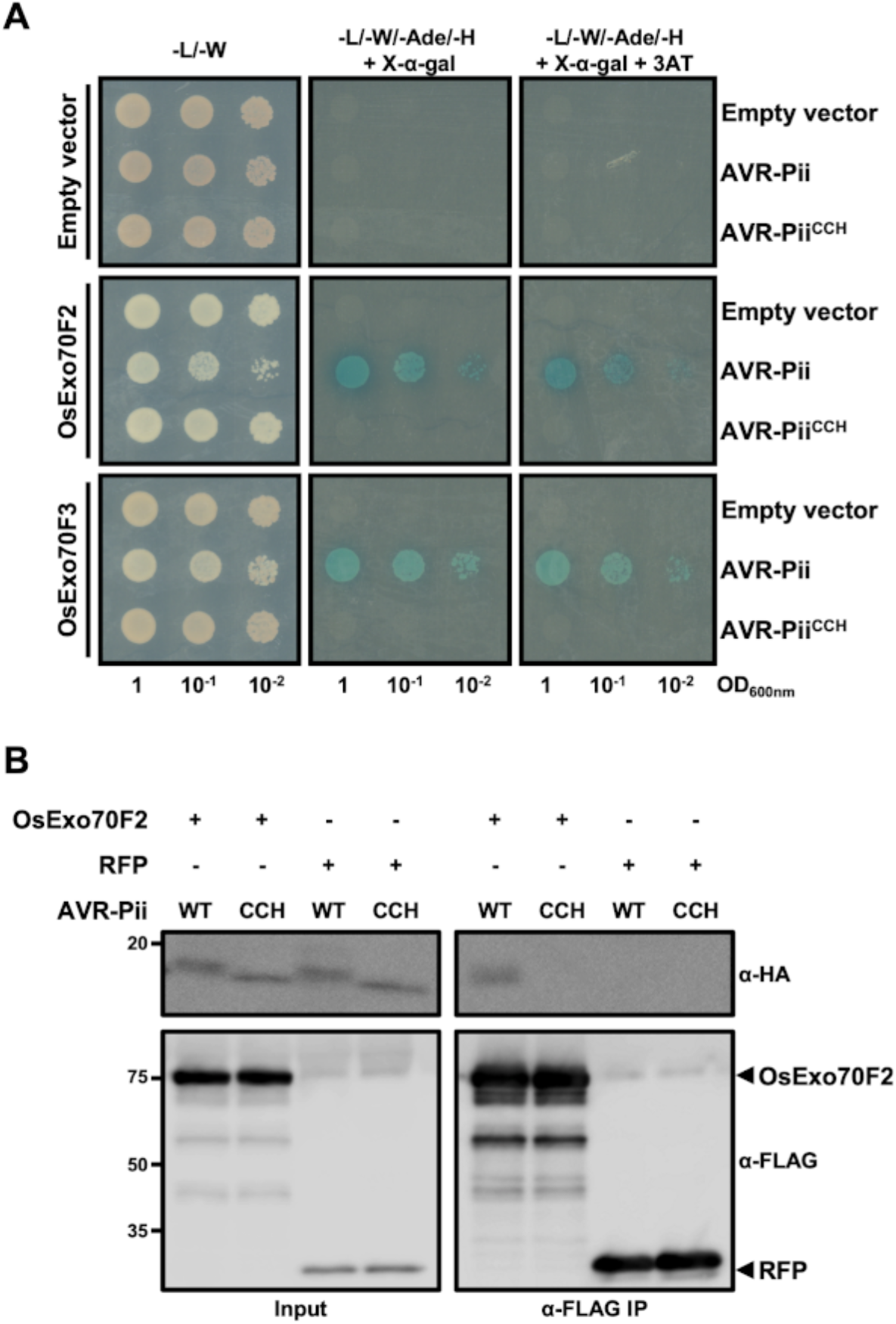
Mutations in AVR-Pii Zinc-finger motif abrogate binding to Exo70 host target. **(A)** Yeast two-hybrid assay of AVR-Pii and AVR-Pii^CCH^ with OsExo70F2 or OsExo70F3 host targets. For each combination, 5µl of yeast were spotted and incubated in double dropout plate for yeast growth control (left) and quadruple dropout media supplemented with X-α-gal and 3AT (right). Growth, and development of blue colouration, in the selection plate are both indicative of protein:protein interaction. OsExo70 proteins were fused to the GAL4 DNA binding domain and AVR-Pii to the GAL4 activator domain. Empty vectors were used as negative control in each combination. **(B)** Co-immunoprecipitation of AVR-Pii and AVR-Pii^CCH^ with OsExo70F2. N-terminally SH-tagged AVR-Pii effectors were transiently co-expressed with N-terminally FLAG-tagged OsExo70F2 (left) or RFP (right) in *N. benthamiana*. Immunoprecipitates (IPs) were obtained with M2 anti-FLAG resin and total protein extracts were probed with appropriate antisera.

As previously reported, co-expression of wild-type AVR-Pii with rice OsExo70F2 or OsExo70F3 supports yeast growth on selective media and development of blue coloration as readouts of protein-protein interactions **(Figure 1a)** (De la Concepcion et al., 2022; Fujisaki et al., 2015). By contrast, co-expression with AVR-Pii^CCH^ did not induce growth or development of blue coloration, indicating that these mutations affect effector binding to the host target **(Figure 1a)**. Further, accumulation of the AVR-Pii^CCH^ protein was consistently lower in yeast cells, when compared with the wild-type effector, which may account for some of the differences observed **(Figure S1)**. We tested whether mutations in the Zinc-finger motif of AVR-Pii also alters protein accumulation in plant cells. For this, we transiently expressed an N-terminally FLAG tagged version of AVR-Pii^CCH^ in *Nicotiana benthamiana* and rice protoplasts. Compared to wild-type AVR-Pii, the stability of the Zinc-finger mutant was also compromised in plant cells **(Figure S2)**, further supporting the role of this motif in maintaining an appropriate protein fold.

To discern whether disruption of the AVR-Pii Zinc-finger motif affects binding to OsExo70 targets beyond reduced effector accumulation, we tested the association of AVR-Pii^CCH^ to OsExo70F2 by Co-IP **(Figure 1b)**. For this, we equilibrated the loading of AVR-Pii and AVR-Pii^CCH^ to ensure similar protein amounts in the input. While AVR-Pii successfully associated with OsExo70F2, AVR-Pii^CCH^ was not able to associate **(Figure 1b)**, confirming the results obtained in the Y2H assay **(Figure 1a)**.

Together, we confirmed the importance of the Zinc-finger motif for AVR-Pii function, showing that mutations in the zinc-binding motif compromise protein stability and abrogate binding to effector targets, most likely due to disruption of the protein fold.

### An HMM-based search reveals a family of Zinc-finger (ZiF) fold effectors with similarity to AVR-Pii in *Magnaporthe*

Genome-wide searches for putative effectors, and their grouping into families, is facilitated by structural information (de Guillen et al., 2015; Lazar et al., 2022; Le Naour-Vernet et al., 2023; Seong and Krasileva, 2021; Win et al., 2012). Given the importance of AVR-Pii residues in forming the ZiF fold, and consequently for function and stability, we reasoned they would be a conserved core maintained during evolution and could be a signature to search and identify related blast effectors.

We created a hidden Markov model (HMM) representing the ZiF fold from *M. oryzae* sequences closely related to AVR-Pii and performed an HMM search against reported proteomes of *M. oryzae* (Chiapello et al., 2015; Yoshida et al., 2016). After filtering out redundant sequences and proteins without effector characteristics, such as the presence of a secretion peptide, we revealed 33 effectors containing a putative Zinc-finger fold that could be classified in 10 different tribes ranging in size from 68 to 119 residues **(Figure 2a, Figure S3).** AVR-Pii belongs to tribe VIII, which contains 7 different alleles **(Figure S3)**. While these ZiF effector tribes were highly divergent at the sequence level **(Figure S3)**, structure prediction by Alphafold2 identified the presence of a ZiF fold in all the tribes **(Figure 2b and c).**

**Figure 2.**
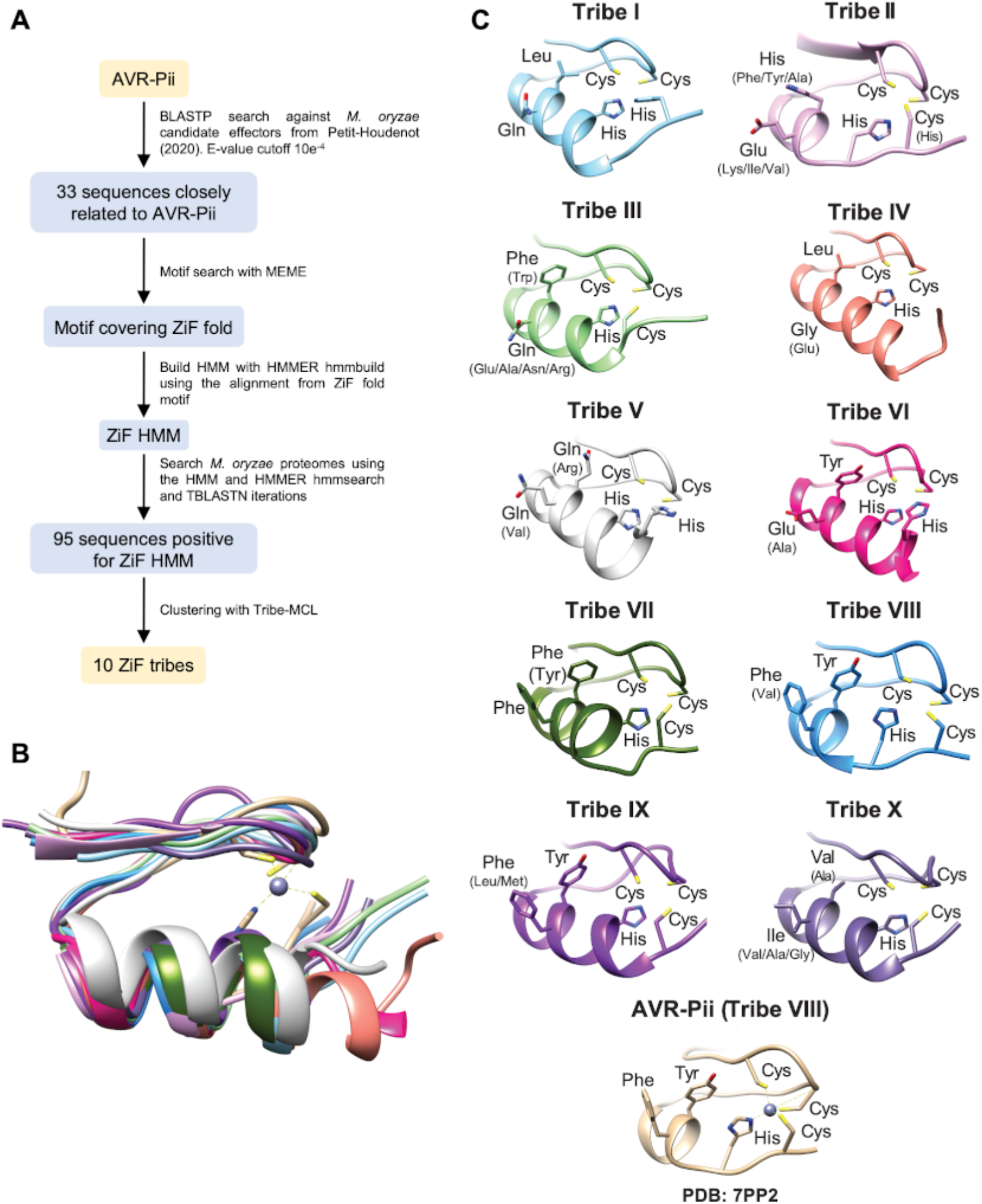
AVR-Pii Zinc-finger Fold (ZiF) defines an effector family in *M. oryzae*. **(A)** Workflow of HMM-based search for Zinc-finger fold effectors. Schematic representation of the ZiF motif search pipeline and results. **(B)** Superimposition of the ZIF fold of AVR-Pii (light brown) and representative members of all ZIF effector tribes of *M. oryzae*. **(C)** Alphafold2 models of amino acid residues that form the binding interface and the zinc finger motif. Amino acid variation within tribes is shown in parentheses. The experimental model of AVR-Pii (PDB: 7PP2) is added as a reference.

### Zinc-finger fold effectors are widespread in host-specialized genetic lineages of the blast fungus

AVR-Pii presents patterns of presence/absence in *Magnaporthe* spp. genomes and was reported to be lost in most rice blast isolates (Latorre et al., 2020; Yoshida et al., 2016). Therefore, we investigated the prevalence and diversity of the ZiF effector family within the genomes of multiple blast isolates with different host specificities. For this, we explored the presence of members of each ZiF effector tribe across 107 *M. oryzae* genome assemblies infecting 13 grass species, as reported previously (Bentham et al., 2021).

We performed iterative TBLASTN searches using a member of each ZiF effector tribe to retrieve sequence related effectors **(Supplemental dataset 1)**. We then mapped the presence/absence of each ZiF effector tribe to every genome of the genetic lineage of *M. oryzae* **(Figure 3)**. For each isolate, we report the absence of a ZiF tribe when TBLASTN did not find a significant hit, and the presence on detection of one or more hits. We also investigated the genomic position of these hits, to consider whether a blast isolate harbours more than one member of a particular ZiF effector tribe. We further define cases of pseudogenized ZiF effectors when the hit from TBLASTN misses a start codon, are truncated, carry an early stop codon, and/or are missing the predicted Zinc-finger motif.

**Figure 3.**
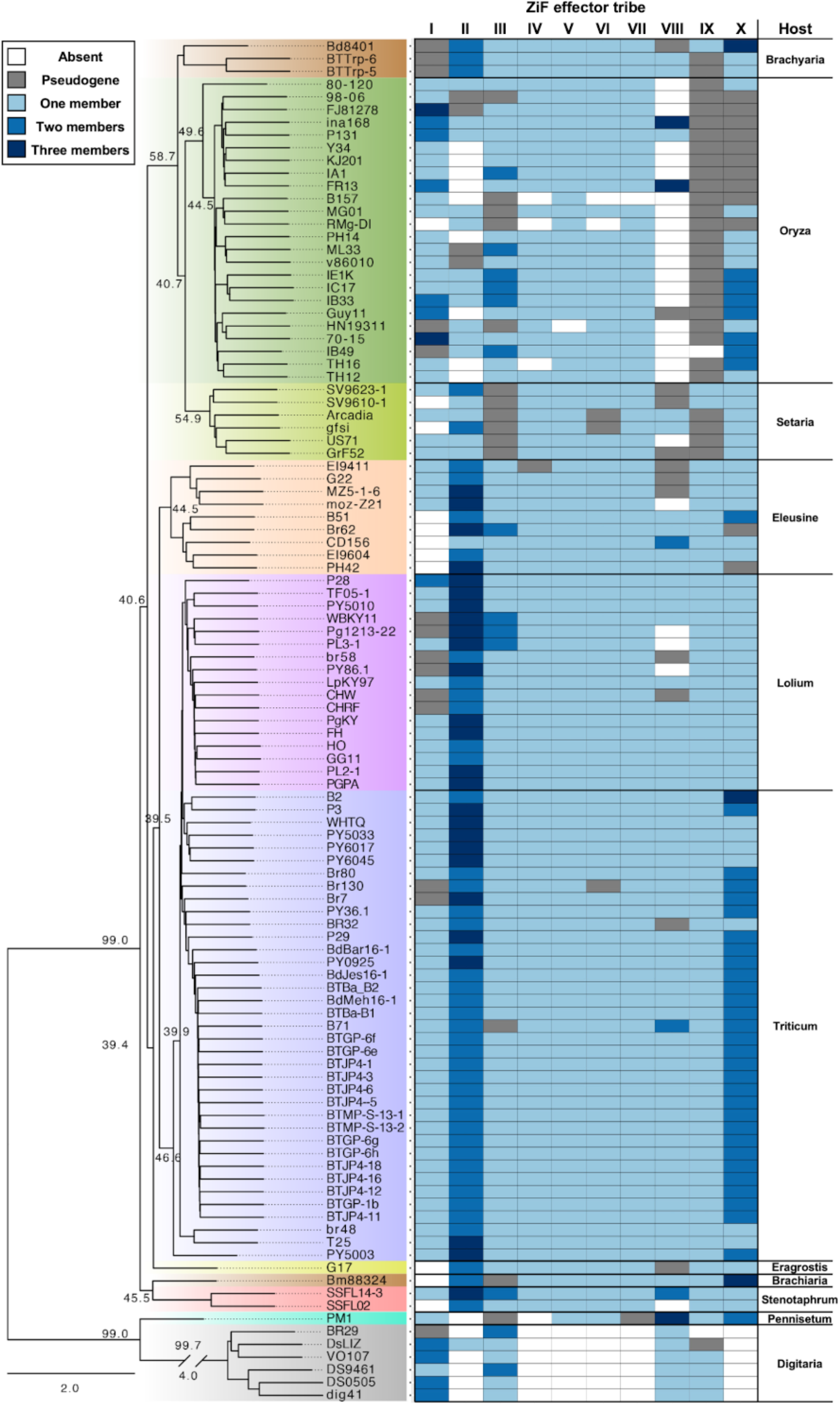
Zif effectors are conserved across different host-specific lineages of *M. oryzae*. Presence/absence analysis of ZiF effector tribes across host-specific lineages of *M. oryzae*. An ASTRAL multispecies coalescence tree (reprinted of a figure panel from Bentham et al. (Bentham et al., 2021)) is shown on the left. On the right, we indicate the presence of ZiF effectors for each tribe in every blast isolate. ZiF tribes without presence in the genome of a blast isolate is represented as white. The presence of a pseudogenized effector (no start codon and/or truncation of the ZiF fold) are represented in grey. When more than one copy of an effector clustering in the same tribe are present, it is represented in different shades of blue.

In total, we identified 95 putative ZiF effector proteins **(Supplemental dataset 2)**. Despite AVR-Pii being reported as absent in most rice infecting blast isolates, our analysis found that most of the ZiF effector family tribes are present in all *M. oryzae* genomes examined **(Figure 3)**. Several of the ZiF effector tribes are also present in the genomes of the more distantly related *Magnaporthe grisea* isolates infecting crabgrass (*Digitaria sanguinalis*) and fountain grass (*Pennisetum americanum*) (**Figure 3)**.

Interestingly, some ZiF effector tribes showed lineage-specific patterns of presence/absence polymorphism **(Figure 3),** which could be associated with colonization of a different host (host jump) (Inoue et al., 2017), or to the genetic conflict with an immune receptor recognizing the effector (Latorre et al., 2020). For example, most rice-infecting *M. oryzae* isolates lack members from the ZiF tribe VIII, which includes AVR-Pii, likely as a consequence of the deployment of the Pii resistance in rice **(Figure 3)**. Furthermore, rice blast isolates did not harbour any member of tribe IX, which may indicate the presence of selective pressure to the pseudogenization of members of this tribe from rice-infecting blast isolates. We also find similar patterns in non-rice infecting *M. oryzae* isolates, such as those infecting foxtail millet (*Setaria spp.*), ryegrass (*Lolium spp*.), signalgrass (*Brachiaria spp.*) and Goosegrass (*Eleusine spp.*) where some specific tribes have been deleted or pseudogenized **(Figure 3)**. Together, this data shows that ZiF effectors form a conserved family in *Magnaporthe* genomes and illustrate the presence of lineage-specific selective pressure that generates specific patterns of effector presence/absence, likely through the genetic conflict with plant immune receptors from different plant species.

### ZiF effectors have different expression patterns during infection

*Magnaporthe* effectors are commonly upregulated during infection (de Guillen et al., 2015; Le Naour-Vernet et al., 2023; Mosquera et al., 2009; Yan et al., 2023). To look for ZiF effector expression during infection, we screened a time-resolved transcriptome of Guy11 infecting the rice cultivar CO39 (Yan et al., 2023) **(Figure S4)**.

Blast strain Guy11 only contains ZiF effectors from the tribe I, III, IV, V, VI, VII and X (it does not harbours a functional copy of AVR-Pii) **(Figure 3)** and we could only detect expression for tribes I, VI and X, meaning that ZiF effector tribes III, IV, V and VII are not expressed (or under the detectable range) during Guy11 infection on CO39 rice cultivars.

ZiF effectors from tribe I and X have their maximum expression peak at 24 and 48 hours after infection (hpi), suggesting a role of these effectors during pathogen colonization **(Figure S4).** By contrast, the effector from ZiF tribe VI is expressed at 96 hpi, which may indicate a role during later stages of the infection **(Figure S4)**.

Both tribe I and X have two members in the genome of Guy11 and they have differences in their expression patterns, suggesting some possible non-overlapping roles of effectors from the same tribe. Both ZiF_If and ZiF_Ik have their expression height at 24 hpi however, ZiF_Ik could be detected at 16hpi but not at 48 hpi while ZiF_If is clearly expressed at 48 hpi but not before 24 hpi **(Figure S4)**. In the case of tribe X this difference is more extreme as we find expression of ZiF_Xp 24 and 48 hpi but we detect no expression of the other member of the tribe, ZiF_Xi **(Figure S4)**.

When considered together, these observations provide evidence that ZiF effectors are co-ordinately expressed during pathogen colonization, and therefore likely to carry out diverse roles during infection.

### An AVR-Pii-like ZiF effector from wheat blast lineages binds Exo70s

Despite being absent in most rice-infecting *M. oryzae* isolates, AVR-Pii-like tribe VIII is highly conserved in wheat blast lineages, including the pandemic B71 clonal lineage (Latorre et al., 2023), and the host Exo70 binding residues identified for AVR-Pii (VIIId) (De la Concepcion et al., 2022) are conserved in the wheat blast counterpart (VIIIc) **(Figure S5)**.

We used Y2H to compare the binding of these two effectors to the Exo70 target. This assay showed that both effectors interact with OsExo70F3, inducing a similar growth of yeast in selective media and development of blue coloration **(Figure 4a)**. To test whether the binding interface previously described for AVR-Pii (De la Concepcion et al., 2022) is functionally conserved in the wheat blast counterpart, we introduced a Phe66Glu mutation in ZiF_VIIIc. This mutation abolished binding to the host target OsExo70F3 in a yeast-2-hybrid assay **(Figure 4a)**, confirming that this effector uses the same surface to bind Exo70 as AVR-Pii (ZiF_VIIId) (De la Concepcion et al., 2022). Both effectors accumulated at similar levels in yeast cells **(Figure S6)**.

**Figure 4.**
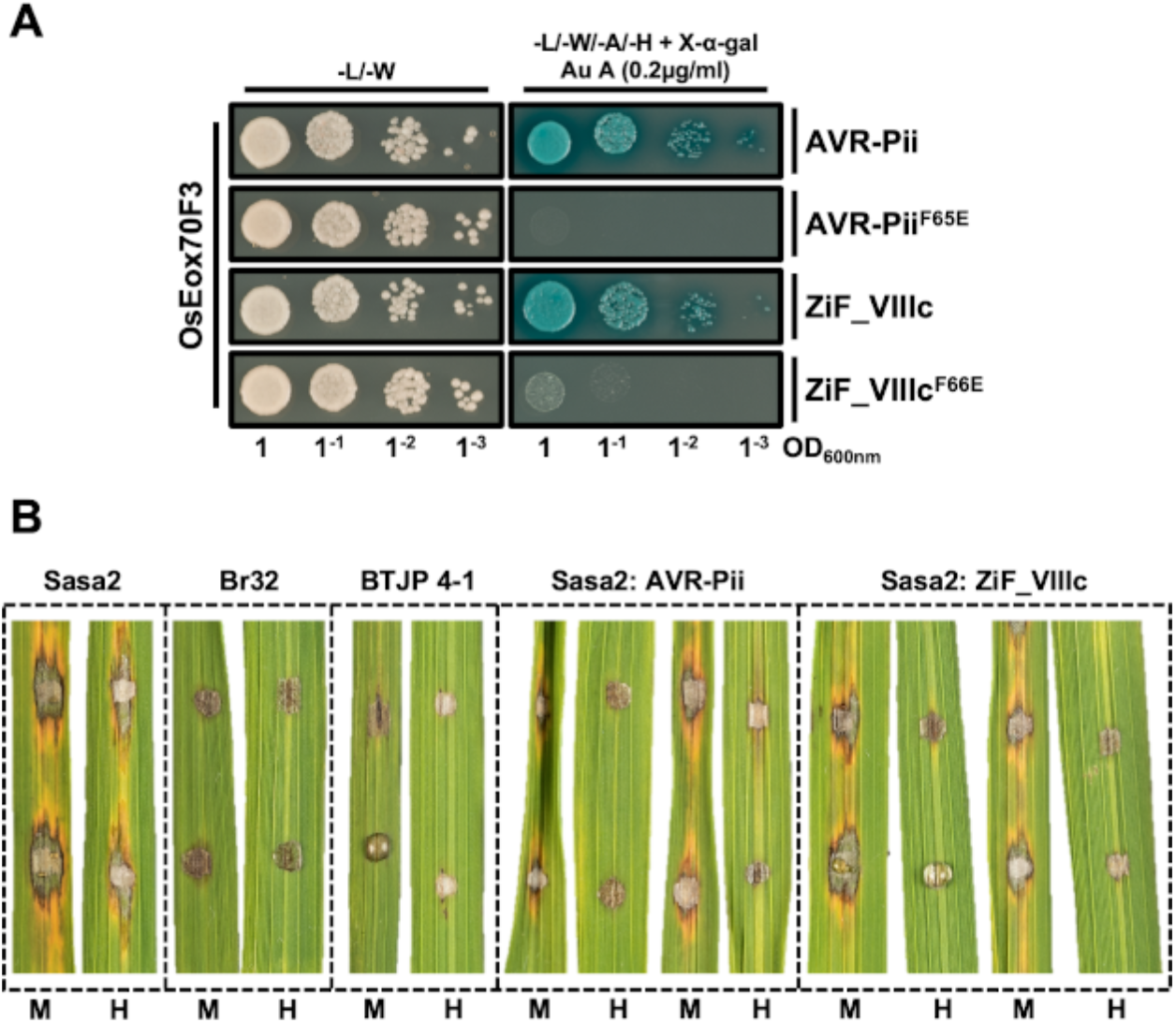
Wheat blast AVR-Pii homolog ZiF_VIIIc binds to OsExo70F3 and is recognized by Pii resistance in rice. **(A)** Y2H assay of wild-type AVR-Pii and ZiF_VIIIc and their corresponding mutants at the host target binding interface (Phe65Glu and Phe66Glu, respectively) to host target OsExo70F3. Left, control plate for yeast growth. Right, quadruple-dropout media supplemented with X-α-gal and aureobasidine A (Au A). Growth and development of blue coloration in the right panel indicates protein-protein interactions. OsExo70F3 was fused to the GAL4 DNA binding domain while effectors were fused to the GAL4 activator domain. Each experiment was repeated a minimum of three times, with similar results. **(B)** Rice leaf blade spot inoculation of transgenic *M. oryzae* Sasa2 isolates expressing AVR-Pii or ZiF_VIIIc from wheat blast isolate BTJP 4-1 in rice cultivars Moukoto (Pii-) and Hitomebore (Pii+). The cultivars Moukoto and Hitomebore are denoted by M and H, respectively. For each experiment, a representative image from replicates with independent *M. oryzae* transformants are shown. Wild-type rice blast isolate Sasa2 and wheat blast Br32 and BTJP 4-1 are included as control. Full images for the three experimental replicates are presented in **Figure S8**.

As ZiF effectors from Tribe VIII are conserved in non-rice infecting isolates, we investigated the presence of potential host Exo70 targets in grasses other than rice. For this, we analysed the conservation of the effector binding interface in recently annotated Exo70F2 and Exo70F3 proteins from eight grass species including rice, wheat, barley and foxtail millet (Holden et al., 2022). This sequence alignment showed that, with the exception of Exo70F2 in Foxtail millet, there is conservation of the hydrophobic pocket where ZiF effectors could likely bind via the docking of a Phe residue, as identified in the AVR-Pii/Exo70F2 complex (De la Concepcion et al., 2022) **(Figure S7)**.

Our data reveal that effectors belonging to the conserved ZiF tribe VIII in *M. oryzae* isolates infecting wheat and other grasses can potentially bind to host Exo70 proteins from clade F in a similar manner as described for the rice blast effector AVR-Pii.

### Zif effectors conserved in wheat blast lineages can be recognized by the rice NLR Pii

ZiF effectors from tribe VIII are conserved in wheat infecting lineages of *M. oryzae*, notably the pandemic clone B71 that is threatening wheat production in three continents (Latorre et al., 2023) harbours at least two copies **(Figure 3)**. Given that ZiF_VIIIc, an effector from Tribe VIII of wheat-infecting isolates, interacts with Exo70 host proteins in a similar way to AVR-Pii **(Figure 4)**, we tested whether this effector triggers Pii-mediated resistance.

For this, we cloned ZiF_VIIIc from the wheat-infecting *M. oryzae* isolate BTJP 4-1 and transformed into the rice-infecting isolate Sasa2, which lacks AVR-Pii, and tested pathogen recognition by rice cultivar Moukoto (Pii-) and Hitomebore (Pii+) **(Figure 4b, Figure S8)**. Spot inoculation assays showed that Sasa2 transformants harbouring the BTJP 4-1 ZiF_VIIIc effector become avirulent on Hitomebore but not on Moukoto **(Figure 4b, Figure S8)**. This recognition is consistent with the result of Sasa2 transformants harbouring AVR-Pii **(Figure 4b, Figure S8)** (De la Concepcion et al., 2022; Fujisaki et al., 2015; Yoshida et al., 2009). We also included in the assay wheat-infecting M. oryzae isolates Br32 and BTJP 4-1 although, as expected, they were unable to infect either Moukoto or Hitomebore plants **(Figure 4b, Figure S8)**.

Together, our data shows that conserved ZiF effectors present in wheat-infecting *M. oryzae* isolates can be recognized by Pii-mediated resistance in rice. This presents new opportunities to explore the transfer of Pii-mediated resistance from rice into other grasses affected by blast disease, including wheat and barley.

### Diverse tribes of ZiF effectors may have different host targets

Although ZiF effectors from different tribes are predicted to share structural similarity **(Figure 2)**, the residues of AVR-Pii that mediate binding to OsExo70 targets are not fully conserved between tribes **(Figure S3)**. We therefore hypothesised that some ZiF effector tribes might target other host proteins.

To test this, we selected the ZiF effector allele from each tribe that is most predominant in the genome of rice-infecting *M. oryzae* lineages and performed a Y2H assay to test for binding to OsExo70F3. ZiF_VIIId (AVR-Pii, the most representative allele of tribe VIII in rice isolates) and AVR-Pii Phe65Glu acted as positive and negative controls, respectively (De la Concepcion et al., 2022). Interestingly, only ZiF_VIIc and ZiF_VIIId interacted with OsExo70F3 in this assay **(Figure 5a)**. All ZiF effectors except ZiF_Xi were produced in yeast cells at similar levels to AVR-Pii (ZiF_VIIId) **(Figure S9)**. Our results showed that ZiF effectors from tribes other than VII and VIII do not interact with OsExo70F3 and may therefore have alternative host targets. The identification and characterisation of alternative host targets will be the subject of future work.

**Figure 5.**
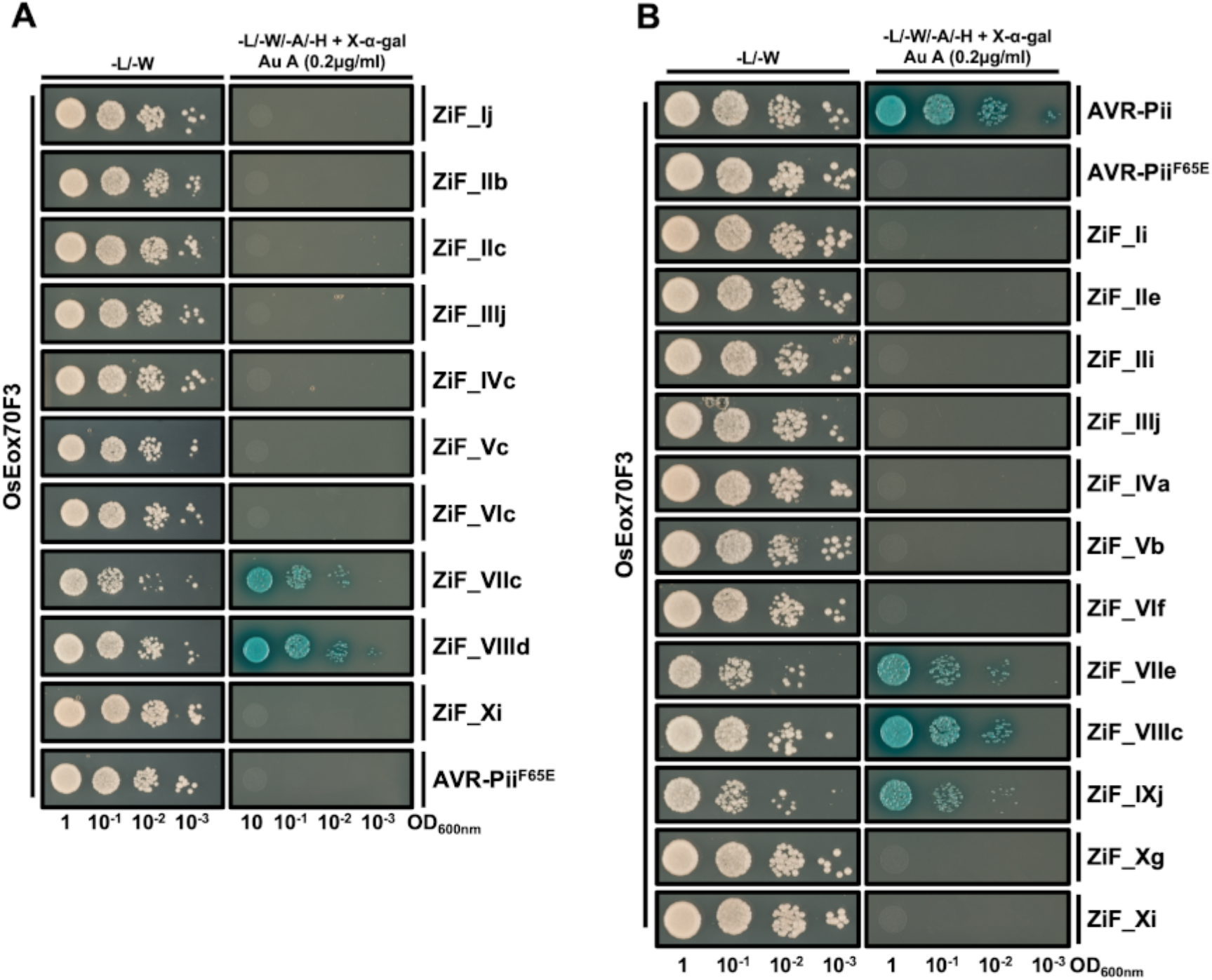
ZiF effectors do not share the same host target. **(A)** Y2H binding assay of ZiF effectors from rice blast isolates to host target OsExo70F3. For each tribe, the most prevalent allele in rice blast lineages was fused to the GAL4 activator domain and co-expressed in yeast cells with OsExo70F3 fused to GAL4 DNA binding domain. AVR-Pii Phe65Glu was used as negative control as previously reported (De la Concepcion et al., 2022). **(B)** Y2H binding assay of ZiF effectors of wheat blast isolates to host target OsExo70F3. The most prevalent effector allele in each ZiF tribe of *M. oryzae* isolates infecting wheat was fused to the GAL4 activator domain and co-expressed in yeast cells with OsExo70F3 fused to GAL4 DNA binding domain. Rice blast AVR-Pii and AVR-Pii Phe65Glu were used as positive and negative controls, respectively. For both assays, a control plate for yeast growth is presented on the left and a plate with quadruple-dropout media supplemented with X-α-gal and aureobasidine A (Au A) is presented on the right. Growth and development of blue coloration in the right panel indicates protein-protein interactions. Each experiment was repeated a minimum of three times, with similar results.

As host-specific *M. oryzae* effector alleles can differ in their association to host interactors (Bentham et al., 2021), we extended the Y2H interaction assay with OsExo70F3 to ZiF effector alleles predominant in *M. oryzae* isolates infecting wheat, which have more representatives of ZiF tribes than rice-infecting isolates **(Figure 3)**. Y2H assays showed that three of these tribes (VII, VIII and IX) interacted with OsExo70F3 **(Figure 5b)**. All wheat blast ZiF effectors, except for ZiF_IVa and ZiF_Xi, had similar protein accumulation in yeast cells **(Figure S10)**.

Together, these Y2H assays indicate functional diversity in the host binding properties of ZiF effector family members, despite their conserved fold.

## Discussion

The study of pathogen effectors and their functions has the potential to generate new biological understanding (Białas et al., 2018) and aid the development of disease management strategies to limit yield losses caused by biotic stress (Cadiou et al., 2023; Zdrzałek et al., 2023). However, the high variability and diversity of effectors at their sequence level, along with their typical small size, makes their classification and functional annotation from sequence challenging (Lovelace et al., 2023; Sperschneider, 2020).

In this study, we showed how the ZiF protein fold observed in AVR-Pii (De la Concepcion et al., 2022) is essential for effector stability, and binding to host targets. We used this fold to define a diverse family of ∼100 secreted proteins, uncovering a previously hidden conservation of these ZiF effectors across *M. oryzae* lineages infecting different grasses and opening new research avenues, both for discovering new effector activities and bioengineering disease resistance.

Previous classification of pathogen effectors into structurally related families based on HMM analysed required pre-existing experimental knowledge of the protein structure (de Guillen et al., 2015; Lazar et al., 2022; Win et al., 2012; Yu et al., 2022). The revolution of protein fold prediction, largely fuelled by the release of AlphaFold2 (Jumper et al., 2021), allowed the structural classification of effectors (Derbyshire and Raffaele, 2023; Seong and Krasileva, 2021; Seong and Krasileva, 2023) and to predict their targets (Homma et al., 2023) without the need of previous structural knowledge. Interestingly, structure prediction approaches did not identify the ZiF effector family in *M. oryzae* (Derbyshire and Raffaele, 2023; Seong and Krasileva, 2021; Seong and Krasileva, 2023), showing how this approach can have limitations when it comes to small, unstructured protein sequences as often is the case of pathogen effectors.

For example, the fold of ∼8000 from over 26000 secreted proteins could not be successfully predicted by AlphaFold2 in a large structural modelling study of secreted proteins from phytopathogenic fungi, including *M. oryzae* (Seong and Krasileva, 2023). Moreover, over 30% of *M. oryzae* secreted proteins could not be predicted by TrRosetta (Seong and Krasileva, 2021) or AlphaFold2 (Seong and Krasileva, 2023).

With less than 100 residues and a large unstructured region at the N-terminus (De la Concepcion et al., 2022), AVR-Pii and other ZiF effectors are likely below the cutoff of structural prediction approaches. However, the previously existing structure of AVR-Pii (De la Concepcion et al., 2022) could still successfully guide HMM searches to identify the ZiF family, illustrating how experimental knowledge of effector structures is still an important foundation for discoveries on effector biology (Mukhi et al., 2020).

Despite AVR-Pii being absent in most rice-infecting *M. oryzae* lineages (Latorre et al., 2020; Yoshida et al., 2009; Yoshida et al., 2016), our combined MEME/HMM/TBLASTN approach uncovered a conservation of the ZiF effector family across all *M. oryzae* isolates. Some of the 10 ZiF tribes identified in this study display presence/absence patterns in some *M. oryzae* lineages, particularly those infecting rice. Interestingly, wheat-infecting isolates conserve all ZiF effector tribes, possibly due to the recent expansion of these lineages (Latorre et al., 2023) causing them to be less polymorphic in comparison to rice-infecting isolates.

An interesting case is the ZiF tribe VIII (that includes AVR-Pii), mostly absent in rice infecting isolates likely as a consequence of the deployment of the Pii resistance in rice. In contrast, all wheat-infecting isolates analysed here includes a member of this effector tribe. This includes the pandemic B71 clonal lineage that has spread and caused disease in three continents (Latorre et al., 2023). We showed how ZiF_VIIIc from these lineages binds and triggers Pii resistance, opening the possibility of transferring Pii resistance from rice to wheat via genetic engineering to rapidly deploy resistance against pandemic wheat blast strains.

We further identify two other ZiF effector tribes (VII and IX) that bind to OsExo70F3. Intriguingly, tribe VII effectors were found to not be expressed during rice infection experiments with rice-infecting isolate Guy11 **(Figure S4)**, and tribe IX members are pseudogenes in the genomes of rice-infecting isolates **(Figure 3)**. Future work could test whether silencing or deletion of these effectors from the genome arose in these isolates to evade activation of Pii or any other resistance genes in rice. A recent study tested this hypothesis for *M. oryzae* MAX fold effectors that are deleted in the genomes of rice blast isolates, however, the authors could not validate that these effectors induce non-host resistance in rice (Le Naour-Vernet et al., 2023).

Other putative effectors following presence/absence distributions are strong candidates for being recognised by a cognate resistance gene. For example, ZiF tribe II, which is absent in multiple rice-infecting lineages is duplicated in wheat-infecting *M. oryzae*, suggesting a resistance gene is present in rice that is unlikely to be *Pii* as the ZiF tribe II effector tested did not interact with OsExo70F3. Similarly, the absence of ZiF tribes in *M. oryzae* isolates infecting wild grasses, such as ZiF tribe I in *Brachyaria-* and *Eleusine*-infecting isolates, highlights the potential of finding new sources of resistance that could be deployed in cultivated grasses.

Further, grass genomes encode NLR proteins with integrated Exo70 domains (Bailey et al., 2018; Brabham et al., 2018). By increasing our knowledge of effectors that bind host Exo70 proteins, we have the potential to engineer Exo70 domains integrated in NLRs to bind and trigger immune responses against these effectors. A similar strategy has been successfully used for MAX effectors binding to HMA integrated domains (Bentham et al., 2022; Cesari et al., 2022; De la Concepcion et al., 2019; Kourelis et al., 2023; Liu et al., 2021; Maidment et al., 2023).

Although all ZiF effector tribes share the same fold, this is not necessarily predictive of protein function but may be involved in maintaining protein integrity while supporting sequence and functional diversification (Bentham et al., 2021; de Guillen et al., 2015). ZiF effectors have expanded in *M. oryzae* lineages and distribute in 10 tribes compared to *M. grisea* lineages, which present only four. This expansion likely reflects a diversification in effector function. Two lines of evidence support of this possibility: First, a previous study reported that the ZiF effector ZiF_IVc (named MoHTR2) is nuclear localized and binds DNA for host transcriptional reprogramming (Kim et al., 2020), a function markedly different from the potential Exo70-binding related function of AVR-Pii. Second, we found that most ZiF effector tribes did not interact with the AVR-Pii host target, OsExo70F3, suggesting alternative targets. Future interaction screens are required to investigate the extent of targets bound by ZiF effectors, potentially generating new fundamental knowledge on *M. oryzae* virulence mechanisms and helping on guiding classical breeding of disease resistance against rice and wheat blast.

## Conclusion

In conclusion, we uncovered a family of effectors with structural similarity to the well-known *M. oryzae* effector AVR-Pii. This family clusters in 10 tribes with a shared fold based on a Zinc-finger motif. Although AVR-Pii is mostly absent in rice-infecting *M. oryzae* lineages, ZiF effector tribes are prevalent in most *M. oryzae* lineages that infect different hosts. We showed that not all ZiF effector tribes share the same target and that AVR-Pii homologues from other lineages can be detected by Pii-mediated resistance from rice. This establishes the principle for investigating whether resistance deployed in rice could be transferred to other grasses as a strategy for developing resistance against pandemic fungal diseases such as wheat blast.

## Materials and Methods

### Cloning of AVR-Pii for expression in planta

For the expression of SH-AVR-Pii in plant cells, we used the expression vector pCAMBIA-SH-AVR-Pii previously described by Fujisaki et al. (Fujisaki et al., 2015). To generate a pCAMBIA-SH-AVR-Pii^CCH^ vector, AVR-Pii^CCH^ cDNA was artificially synthesized by GenScript Japan Inc (Tokyo, Japan) and the coding sequence was amplified by PCR using the primer set: KF1f (5’-AATCACTAGTGGTGGCGGTCTTCCCACTCCGGCCAGCCTG-3’) and KF118r (5’-AATCCTGCAGTTAGTTGCATTTAGCATTAAAATA-3’). Resulting PCR product was introduced into pCAMBIA-SH (Fujisaki et al., 2015) using *Spe*I and *Pst*I sites. Transient expression of SH-AVR-Pii and SH-AVR-Pii^CCH^ mutant in *N. benthamiana* was performed as described previously (Kanzaki et al., 2012).

For transient expression of FLAG tagged AVR-Pii and AVR-Pii^CCH^ in rice protoplasts, we used the expression vectors pAHC-FL-AVR-Pii (Fujisaki et al., 2015) and pAHC-AVR-Pii^CCH^. For the construction of pAHC-FL-AVR-Pii^CCH^, AVR-Pii^CCH^ cDNA was amplified by PCR using primer set; KF1f and KF3r (5’-AATCGGATCCTTAGTTGCATTTAGCATTAAAATA -3’). The resulting PCR product was exchanged with wild type AVR-Pii coding sequence of pAHC-FL-AVR-Pii using *Spe*I and *BamH*I sites.

### Cloning of ZiF for Yeast-2-Hybrid assays

Coding sequences for the effector domain of the ZiF proteins reported here were synthesised as double-stranded DNA fragments (gBlocks, Integrated DNA Technologies). The fragments were subsequently inserted via BsaI in a pGADT7 vector adapted for Golden Gate cloning provided by the SynBio service at The Sainsbury Laboratory, Norwich.

To generate AVR-Pii^CCH^ mutant in pGADT7, AVR-Pii^CCH^ coding sequence was introduced into pGADT7 using EcoRI and BamHI sites following amplification by PCR using the following primer set:

KF777f (5’-CACCGAATTCCTTCCCACTCCGGCCAGC -3’)

KF780r (5’-AACTGGATCCTTAGTTGCATTTAGCATTAAAAT -3’)

Rice blast AVR-Pii and AVR-Pii Phe65Glu in pGADT7, as well as, OsExo70F3 in pGBKT7 were reported previously (De la Concepcion et al., 2022).

### Cloning of ZiF_VIIIc from BTJP 4-1

DNA was extracted from *M. oryzae* isolate BTJP 4-1 using a CTAB method from mycelia grown on PDA plates. The effector AVR-Pii from was obtained via PCR amplification using Phusion Polymerase (New England Biolabs). Primers were designed on the promoter and terminator of ZiF_VIIIc from BTJP 4-1 and included the overhangs required for plasmid assembly. Primer sequences used were:

PCB1532_AVR-Pii_promoter_p1f

(5’-AAGCTGGAGCTCCACCGCGGCGCAGCAGAGGTCTAGTTTGAGCAC-3’)

AVR-Pii_terminator_PCB1532_p1r

(5’-CTATAGGGCGAATTGGGTACCTCGCATCGGCAAATACTCTCGAG-3’)

The amplified fragment including promoter, CDS, and terminator was assembled into the backbone PCB1532 via In-Fusion cloning (In-Fusion cloning kit, Clontech Laboratories).

### Transient expression in rice protoplasts

Rice protoplasts were isolated from rice cell culture (Ichimaru et al., 2022; Ishikawa et al., 2014). For cell wall digestion, 5 ml of packed cell volume of cultured cells was mixed with 25 ml of cellulase solution [4% Cellulase RS (w/v; Yakult, Tokyo, Japan), 4% Cellulase R10 (w/v; Yakult), 0.1% CaCl_2_.6H_2_O (w/v), 0.4M mannitol, 0.1% MES (w/v) pH5.6] and incubated with gentle shaking (60 rpm) at 30°C for 3 h. After filtration through miracloth (Millipore, Billerica, MA), protoplasts in the flow through fraction were collected by centrifugation at 800g and washed two times with 20 ml of W5 buffer (154 mM NaCl, 125 mM CaCl_2_, 5mM KCl and 2mM MES pH5.7). Protoplast concentration was adjusted to 2-3 x 10^6^ protoplasts/ml with W5 buffer. For transfection, 10 µg of plasmids pAHC-FL-AVR-Pii or pAHC-FL-AVR-Pii^CCH^ in a volume of 10 µl was mixed with 100 µl of protoplast solution. Next, 110 µl of PEG solution [40% PEG4000 (w/v; Fluka), 0.2M mannitol, 0.1M CaCl_2_] was added to the protoplast solution, mixed gently, and incubated at room temperature for 20 min. Then, 700 µl of W5 buffer were added, and protoplasts were collected by centrifugation at 800 g. The protoplasts were washed with 700 µl of W5 buffer, resuspended with 500 µl of W5 buffer, and incubated in the dark at 30°C for 20 h. After incubation, protoplasts were collected by 800 g centrifugation, resuspended with 60 µl of GSB buffer [62.5 mM Tris-HCl (pH6.8), 10% Glycerol, 0.2 g/ml SDS, 5µg/ml Bromophenol blue, and 100 mM DTT], and subjected to western blot assay using HRP-conjugated anti-FLAG M2 (Sigma-Aldrich, St. Louis, MO).

### Protein-protein interaction: Yeast-2-hybrid assay

To test the interaction between AVR-Pii^CCH^ and OsExo70F3, Yeast-2-hybrid assays were performed as described previously (Kanzaki et al., 2012). Co-transformed yeast cells were prepared in a dilution series with OD_600_ = 3.0 (x1), 0.3 (×10^-1^) and 0.03 (×10^-2^)] and spotted onto quadruple dropout medium (QDO); basal medium lacking Trp, Leu, Ade and His but containing 5-Bromo-4-Chloro-3-indolyl a-D-galactopyranoside (X-a-gal) (Clontech). To detect interactions, both QDO medium with and without 10 mM 3-amino-1,2,4-triazole (3AT) (Sigma) was used. Yeast cells were also spotted onto double dropout medium (DDO); basal medium lacking Trp, Leu to test cell viability.

To detect protein accumulation in yeast cells, cells were propagated in liquid DDO at 30°C overnight. Forty mg of yeast cells were collected and resuspended with 160 ul GTN + DC buffer [10% glycerol, 25mM Tris-HCl (pH 7.5), 150mM NaCl, 1 mM DTT and 1 tablet of complete EDTA-free (Roche, Basel Switzerland)]. Then, 160 ul of 0.6 N NaOH was added, mixed gently, and incubated at room temperature for 10 min. Next, 160 ul of gel sample buffer [40%(w/v) glycerol, 240 mM Tris-HCl pH 6.8, 8% (w/v) SDS, 0.04% (w/v) bromophenol blue, 400 mM DTT] was added and incubated at 95 °C for 5 min. After centrifugation at 20,000 g for 5 min, the supernatant was subjected to SDS-PAGE. Proteins expressed from bait and prey vectors were detected by using anti-Myc-tag mAb-HRP-DirectT (MBL, Nagoya, Japan) and anti-HA-Peroxidase 3F10 (Roche), respectively.

Yeast-two-hybrid assays of the interaction between ZiF effectors and the host target OsExo70F3 was performed as described previously (De la Concepcion et al., 2019; De la Concepcion et al., 2018; De la Concepcion et al., 2022). In brief, pGADT7 plasmids encoding the effector domain of different ZiF were co-transformed into chemically competent Y2HGold cells (Takara Bio, USA) with a pGBKT7 plasmid encoding OsExo70F3 in using a Frozen-EZ Yeast Transformation Kit (Zymo research).

After growing in selection plates, single co-transformants were inoculated in liquid SD-Leu-Trp media overnight at 30 °C. The saturated culture was used to make serial dilutions of OD_600_ 1, 0.1, 0.01, and 0.001 and 5 µl of each dilution was spotted on a SD-Leu-Trp plate as a growth control, and on a SD-Leu-Trp-Ade-His plate containing X-α-gal and supplemented with 0.2 µg/ml Aureobasidin A (Takara Bio, USA). Plates were imaged after incubation for 60–72 hr at 30 °C. Each experiment was repeated a minimum of three times, with similar results.

To assay the accumulation of protein in yeast cells, total yeast extracts were produced by harvesting cells from the liquid media and incubate them for 10 minutes at 95°C after resuspending them in LDS Runblue sample buffer. Samples were then centrifugated and the supernatant was subjected to SDS-PAGE and western blot. The membranes were probed with anti-GAL4 DNA Binding domain (Sigma) antibody for the OsExo70F3 protein in pGBKT7 and with the anti-GAL4 activation domain (Sigma) antibody for AVR-Pii and ZiF effectors in pGADT7.

### Protein-protein interaction: In planta co-immunoprecipitation (co-IP)

To test for association of AVR-Pii and AVR-Pii^CCH^ to OsExo70F2 in planta, Co-immunoprecipitation (co-IP) was performed as described previously (Fujisaki et al., 2015). In brief, FLAG-OsExo70F2, SH-AVR-Pii and SH-AVR-Pii^CCH^ were transiently and separately expressed in *N. benthamiana* leaves. Since the accumulation level of SH-AVR-Pii^CCH^ was low, SH-AVR-Pii and SH-AVR-Pii^CCH^ proteins were concentrated by immunoprecipitation utilizing HA tag. The amounts of both proteins were adjusted to similar level by dilution using *N. benthamiana* leaf lysates in GTN+DC buffer [10% glycerol, 25mM Tris-HCl (pH 7.5), 150mM NaCl, 1 mM DTT and 1 tablet of complete EDTA-free (Roche, Basel Switzerland)]. Then, the SH-AVR-Pii and SH-AVR-Pii^CCH^ proteins were mixed with leaf lysates containing FLAG-OsExo70F2 protein, and the mixture was used for co-IP using anti-FLAG M2 resin (Sigma-Aldrich, St. Louis, MO).

### Bioinformatic analyses: HMM search in fungal genomes

To develop a motif profile for the ZiF fold present in AVR-Pii, we first searched a database of candidate *M. oryzae* effectors (Petit-Houdenot et al., 2020) for sequences closely related to AVR-Pii using BLASTP from BLAST+ suite (Camacho et al., 2009). We extracted the sequences showing a match to AVR-Pii sequence with an E-value less than or equal to 0.0001. Next, we analysed these sequences using MEME (Bailey et al., 2015) to identify a motif covering the amino acids involved in the ZiF fold. We used this ZiF fold motif alignment from MEME as an input to hmmbuild program from HMMER software (HMMER3/f 3.1b1, http://hmmer.org, May 2013) for creating a hidden Markov model (HMM) that represents the ZiF fold. We did motif searches against the proteomes of *M. oryzae* reported by Chiapello et al. (Chiapello et al., 2015) and Yoshida et al. (Yoshida et al., 2016) using the ZiF fold HMM and hmmsearch program from HMMER. For clustering of ZiF fold motif containing proteins, we used Markov Clustering algorithm (MCL) (Dongen, 2008) with Tribe-MCL option.

### Bioinformatic analyses: TBLASTN search in Magnaporthe genomes

We used a TBLASTN-based approach to identify ZIF-effector members in 107 genomes (Bentham et al., 2021) that represent Magnaporthe isolates collected from 13 host plant species. We used each putative ZIF effector identified by HMM as individual queries for TBLASTN (e-value threshold of 1e-10) and extracted all hits with >80% query coverage. We then deduplicated the hits to generate a non-redundant list of ZIF effector variants and manually curated the hits to identify pseudogenes. This led to the identification of 95 putative ZIF effectors. Alignments of all ZIF tribes were generated using Clustal Omega (Sievers and Higgins, 2014).

### Bioinformatic analyses: Alphafold2 prediction and structural comparison

To predict the structure of ZIF effector candidates, we first removed the signal peptide sequence of the 95 ZIF effector candidate. The processed sequences were then batch submitted to a local installation of Alphafold2 (Jumper et al., 2021) using the default parameters. We then extracted the file ranked_model_0.pdb of each ZIF effector for structural comparison. Automated structural comparison by TM-align failed because ZIF effectors contain a large, unstructured region at their N-terminus. We thus selected representative members of each tribe and superimposed them using ChimeraX (Pettersen et al., 2021) for structural comparison. To avoid errors during superimposition, we removed the unstructured region and used only the structured part of the protein (as shown in Figure 2).

### *M. oryzae* protoplast generation and transformation

Protoplast generation and transformation of *M. oryzae* isolate Sasa2 was carried out as described in Saitoh et al. (Saitoh et al., 2012). Briefly, protoplasts were generated by inoculating 200 ml of Yeast Growth media (5g of yeast extract, 20g of glucose in 1L dH_2_O) with ∼10cm^2^ of mycelia grown on PDA plates and shaking at 121 rpm at ∼24°C for 3 days. Mycelia was harvested via filtration and added to 15 ml lysing enzyme solution (from *Trichoderma harzianum*; Sigma-Aldrich) in a 50ml tube. The tube was shaken at 45 rpm at 30°C for 4 hours. Resulting protoplast suspension was filtered through two layers of Miracloth and collected in a clean tube. An equal volume of 0.6 M STC was added to the filtrate and centrifuged at 2,500 rpm at room temperature for 10 minutes and the supernatant discarded. Protoplasts were gently resuspended in 5 ml of 1 M STC. Additional resuspension and centrifugation steps were repeated with 30 ml, 25 ml, and 5 ml 1M STC sequentially, with the supernatant discarded between each wash step. Protoplasts were adjusted to a final concentration of 1 x 10^8^ cells/ml.

For transformation, 60% PEG solution was added at a volume of 4x the volume of the protoplast suspension. Plasmid DNA solution (4μg of DNA in 0.2ml of H_2_O) was added to 400 μl of protoplast suspension and incubated on ice for 30 minutes. Sequential additions of 60% PEG solution were made of 0.2 ml, 0.3 ml, 0.6 ml, and 1.2 ml with gentle mixing of the solution. The resulting solution was left at room temperature for 15 minutes then centrifuged at 2,500 rpm for 10 minutes and the supernatant discarded. The protoplasts were gently resuspended in 0.3 ml 1 M STC and the final suspension spread onto selective media containing sulfonylurea. Transformants were recovered for 2 days at ∼24°C and monoconidial isolations collected.

Successful transformants were genotyped with AVR-Pii primers BTJP41_AVRPii_t3_p1f 5’ – GCCTCTTCCTGTTCGCCA - 3’ and BTJP41_AVRPii_t3_p1r 5’ – GTGGGAAGGGCTGCGATT - 3’. No amplification was observed from the Sasa2 wild type.

### Rice blast infection assay

To evaluate the response of rice carrying Pii to rice blast fungal strain Sasa2 transformed with ZiF effectors from wheat blast lineages, we performed leaf drop infection assays in detached leaves from rice line Hitomebore (Pii+) and control susceptible line Moukoto (Pii-). Leaves were obtained from three-week old plants and placed on 1% agarose on square petri dishes.

*Magnaporthe* spores were collected from 10 days old cultures of the corresponding strain and diluted to a final concentration of 1×10^5^ conidia mL^-1^ in 0.2% gelatine. Leaves were inoculated with 20 µL of the conidia suspension with two leaves per treatment and 3 biological replicates. Lesions were imaged at 5 days post infection.

---

**Figure S1.**
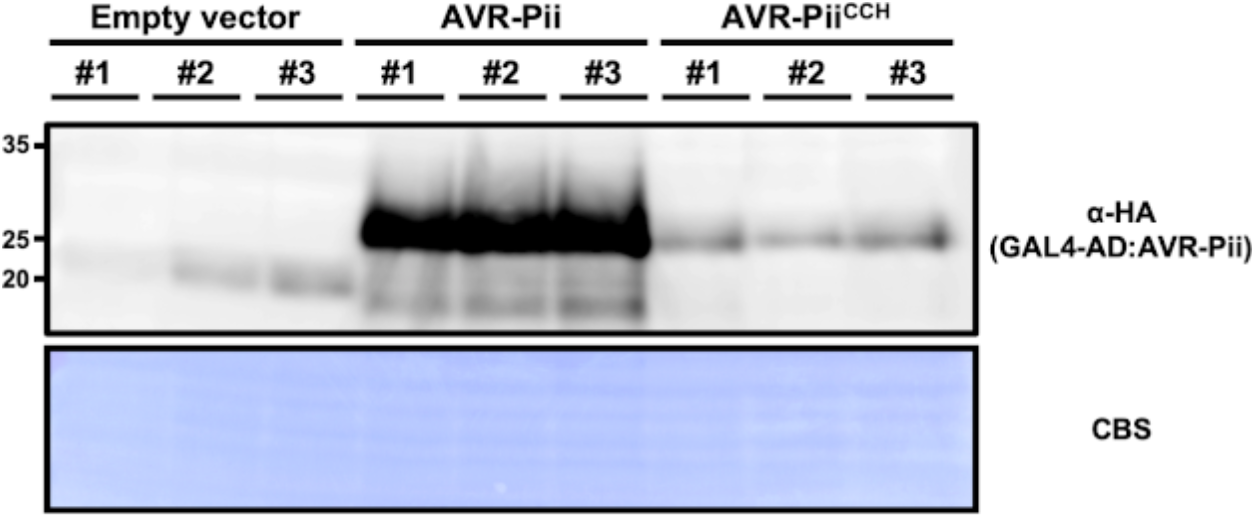
Accumulation of AVR-Pii^CCH^ is compromised in yeast cells. Yeast lysate was probed for the expression of AVR-Pii effectors using anti-HA antibodies for the effectors fused to the GAL4 activation domain (AD). Total protein extracts were coloured with Coomassie Blue Stain (CBS).

**Figure S2.**
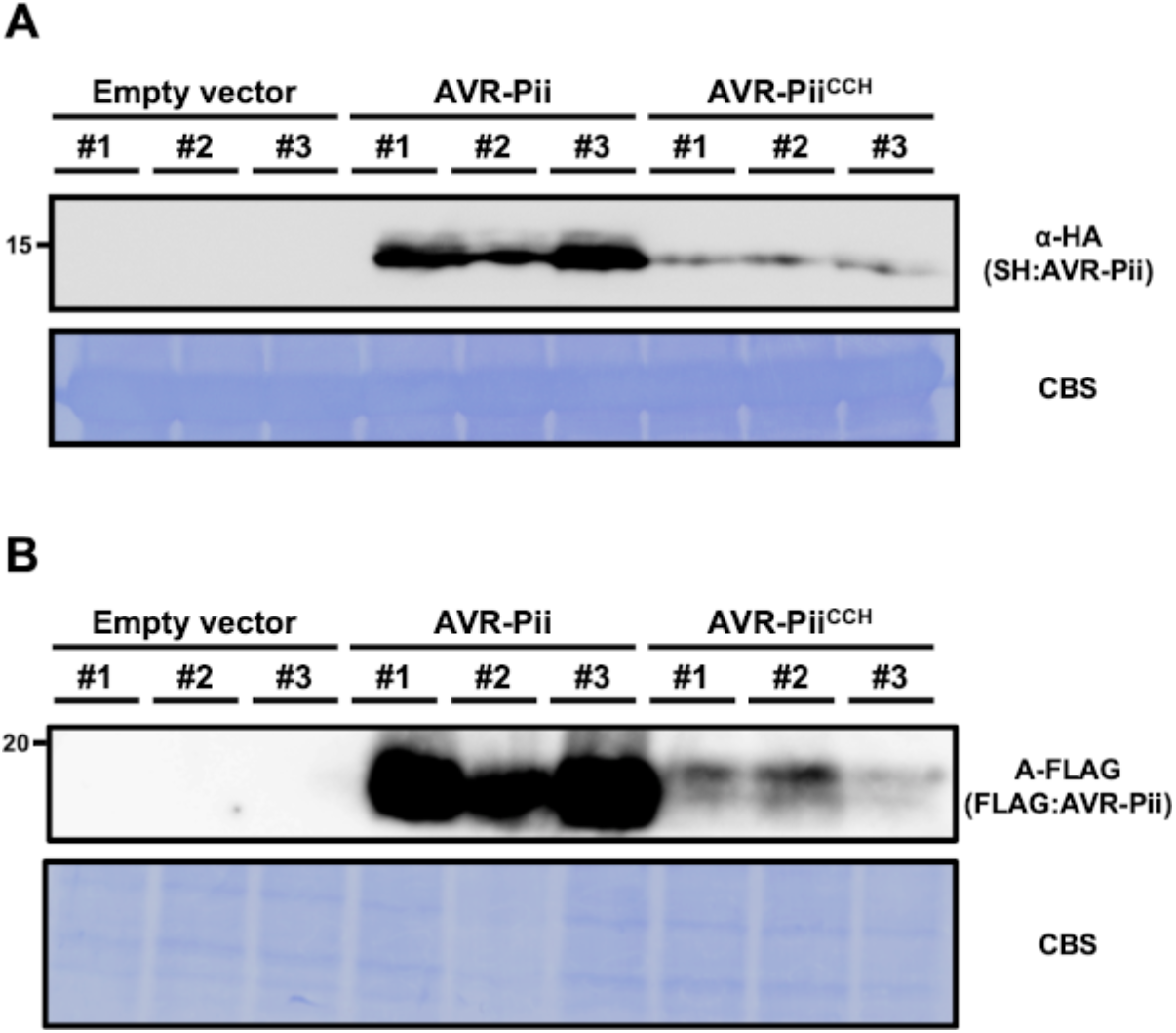
Accumulation of AVR-Pii^CCH^ is compromised in plant cells. **(A)** Western blot analysis of N-terminally SH-tagged AVR-Pii or AVR-Pii^CCH^ transiently expressed in *N. benthamiana*. Plant lysates were probed with anti-HA antibodies for the presence of effectors. Total protein extracts were coloured with Coomassie Blue Stain (CBS). **(B)** Western blot analysis of N-terminally FLAG-tagged AVR-Pii or AVR-Pii^CCH^ transiently expressed in rice protoplasts. Lysates from rice protoplasts were probed with anti-FLAG antibodies for the presence of AVR-Pii effectors. Total protein extracts were coloured with Coomassie Blue Stain (CBS).

**Figure S3.**
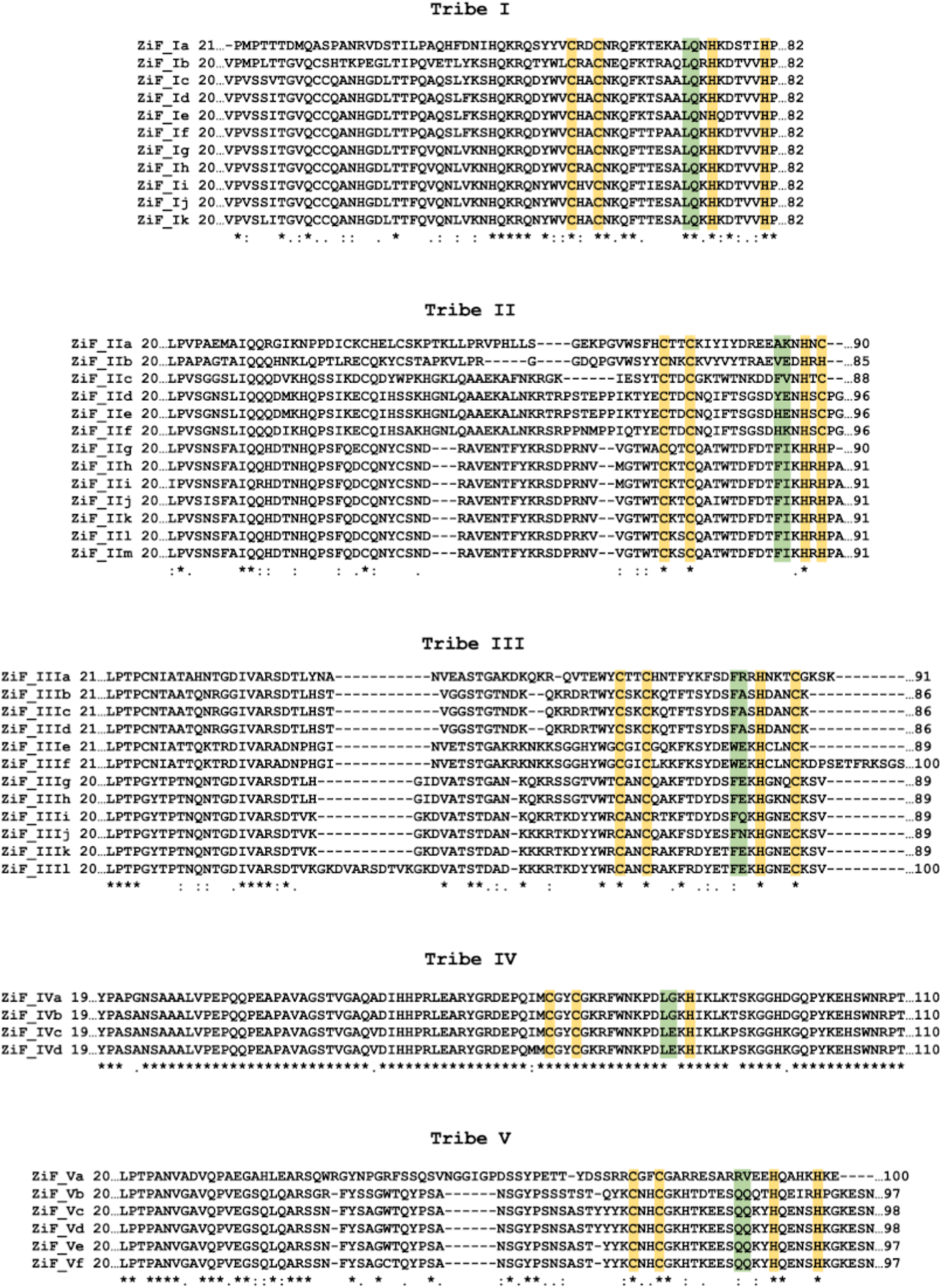
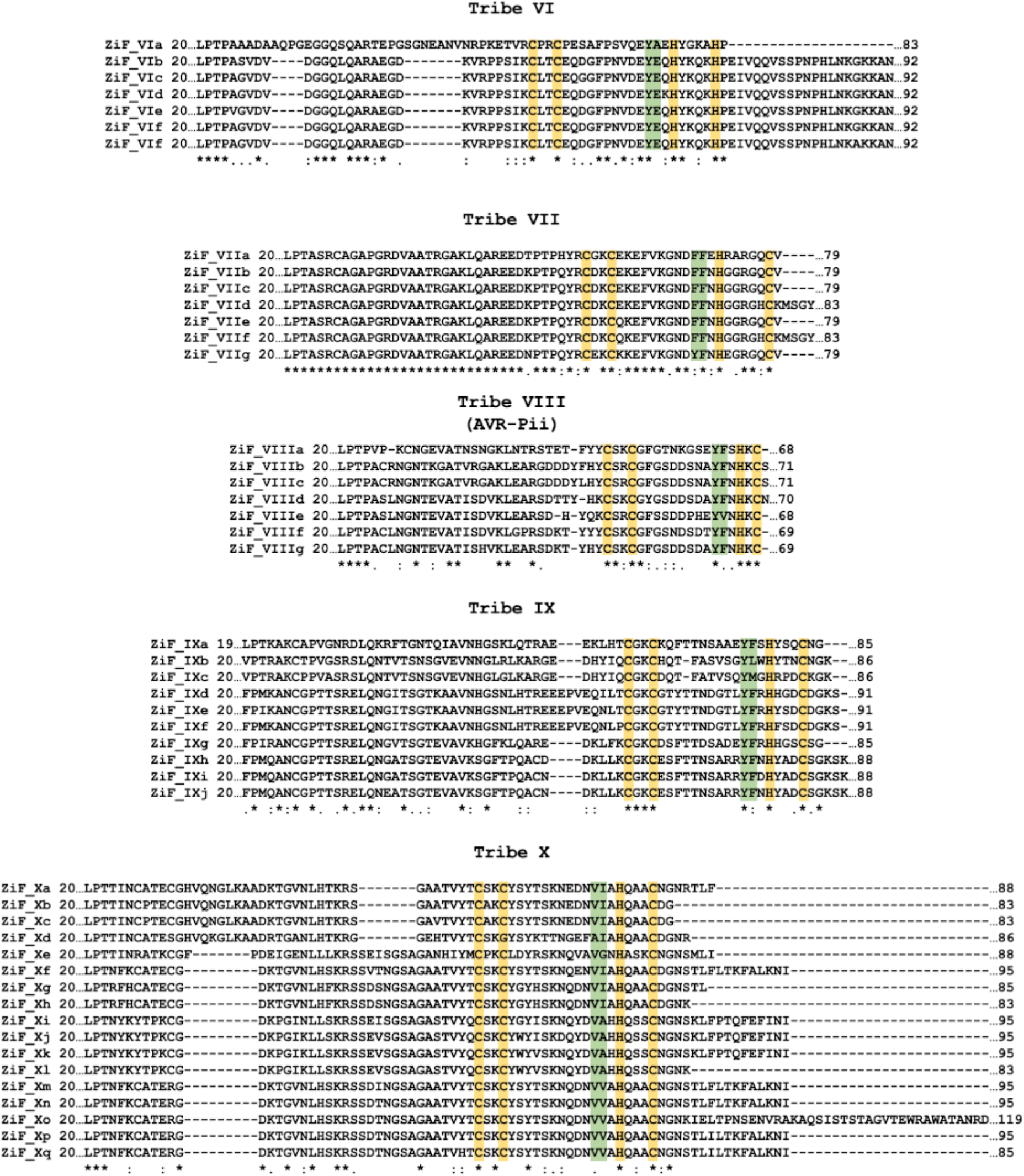
Alignment of ZiF effector tribes showing conservation of the Zinc-finger motif and differences in residues forming the Exo70 binding interface. Protein sequence alignment of ZiF effector proteins reported here separated by tribes generated with Clustal Omega (Sievers and Higgins, 2014). Residues contributing to the formation of a Zinc-finger motif are highlighted in yellow and residues at the equivalent positions of AVR-Pii binding interface (De la Concepcion et al., 2022) are highlighted in green.

**Figure S4.**
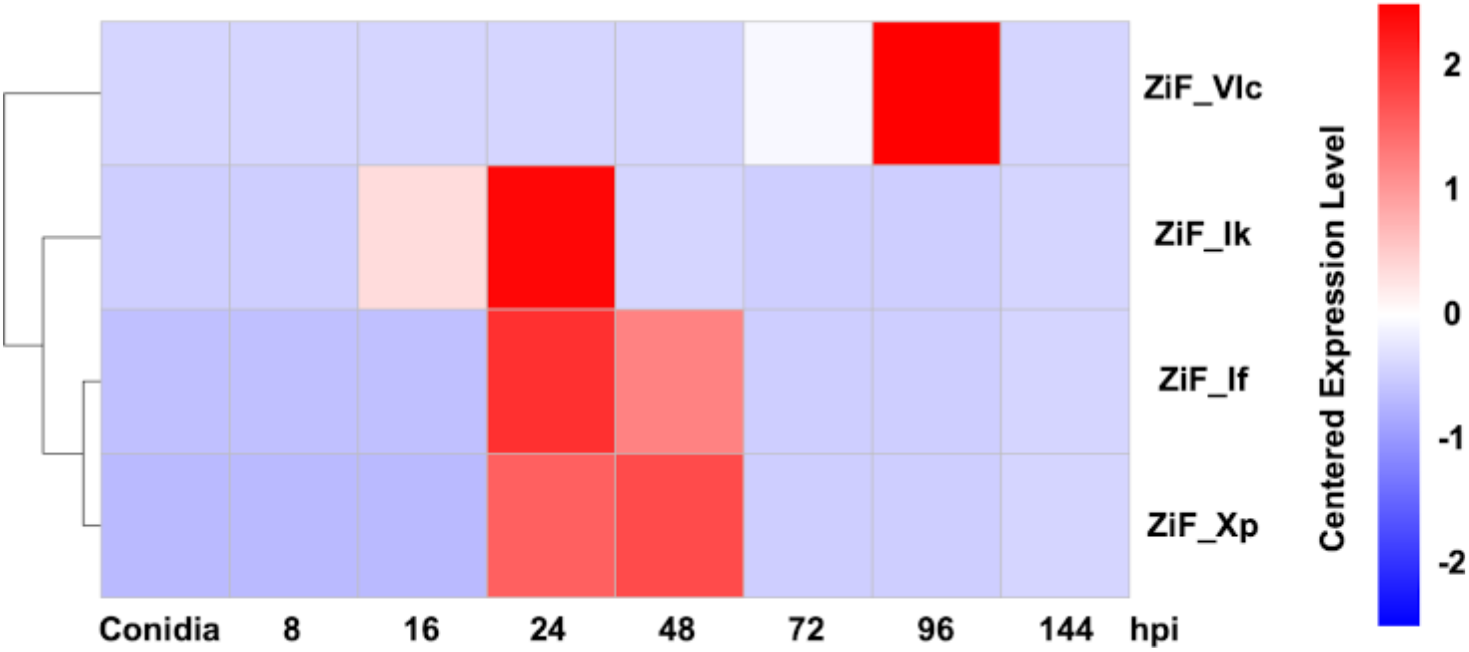
Guy11 ZiF effectors are differentially expressed during infection in rice CO39 cultivar. Heatmap for expression of ZiF effectors at different times after infection (hpi). Expression is represented in shades of red or blue according to the centered expression levels as indicated. Only effectors for which we found expression are represented. Magnaporthe isolate Guy11 does not harbour AVR-Pii (ZiF_VIII).

**Figure S5.**
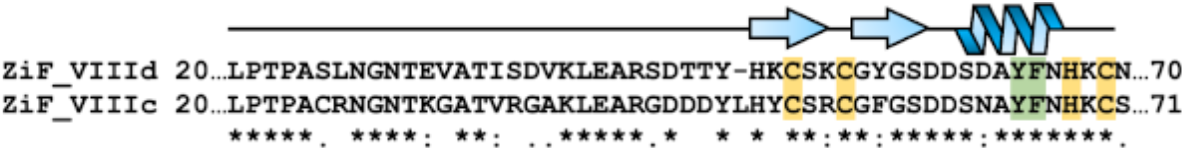
Alignment of ZiF effector tribes showing conservation of rice and wheat blast alleles of AVR-Pii. Protein sequence alignment of ZiF effector proteins ZiF_VIIId (AVR-Pii) and ZiF_VIIIc with Clustal Omega (Sievers and Higgins, 2014). Secondary structure features based on AVR-Pii structure (De la Concepcion et al., 2022) are shown above. Residues contributing to the formation of a Zinc-finger motif are highlighted in yellow whereas the residues at the equivalent positions of AVR-Pii binding interface (De la Concepcion et al., 2022) are highlighted in green.

**Figure S6.**
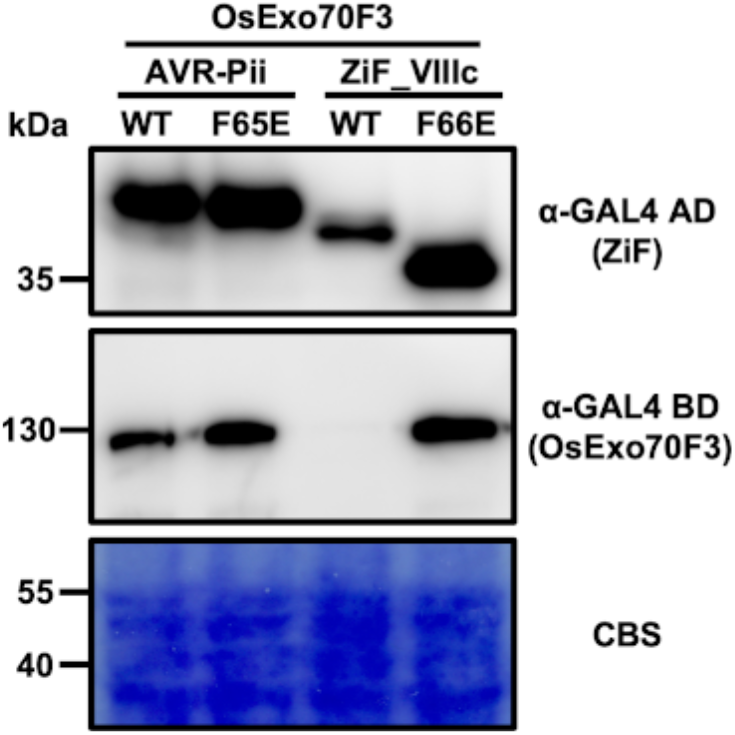
Protein accumulation in Yeast-Two-Hybrid assay analysed by Western blot. Yeast lysate was probed for the presence of OsExo70F3, AVR-Pii, ZiF_VIIIc and their respective mutants using anti-GAL4 binding domain (BD) and anti-GAL4 DNA activation domain (AD) antibodies. Total protein extracts were stained with Coomassie Blue Stain (CBS). OsExo70F3 accumulation is consistently lower in positive interactions with ZiF effectors as noticed here and elsewhere in this and previous studies (De la Concepcion et al., 2022).

**Figure S7.**
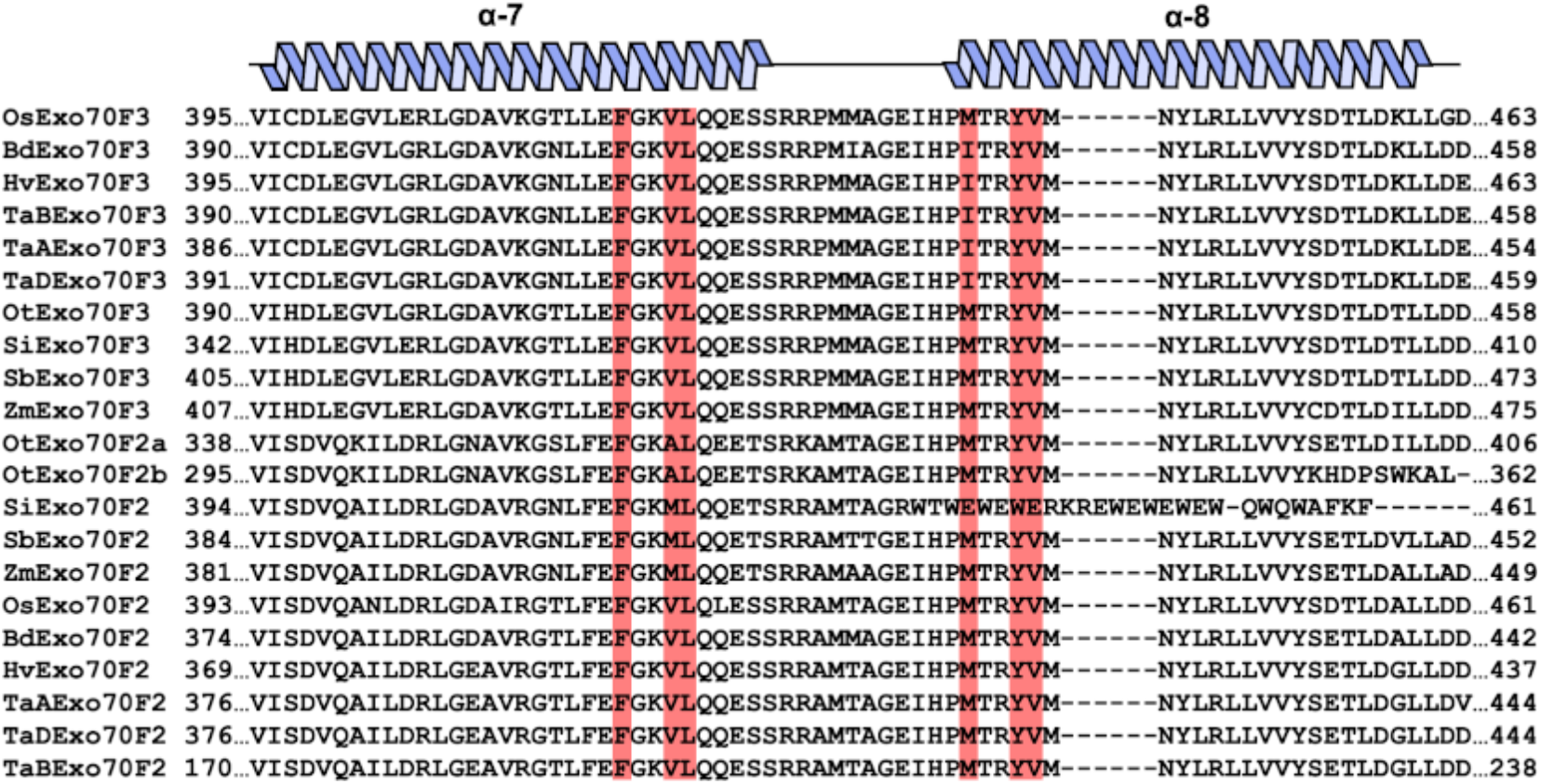
Alignment of grass Exo70 proteins showing conservation at the residues form the OsExo70F3 binding interface with AVR-Pii. Protein sequence alignment for the orthologs of AVR-Pii host targets, OsExo70F2 and OsExo70F3, from different grass species annotated by Holden et al. (Holden et al., 2022). Sequence alignment was generated with Clustal Omega (Sievers and Higgins, 2014). Residues at the equivalent positions of the OsExo70F2 binding interface with AVR-Pii are highlighted in red. Secondary structure features based on OsExo70F2 structure (De la Concepcion et al., 2022) are shown above.

**Figure S8.**
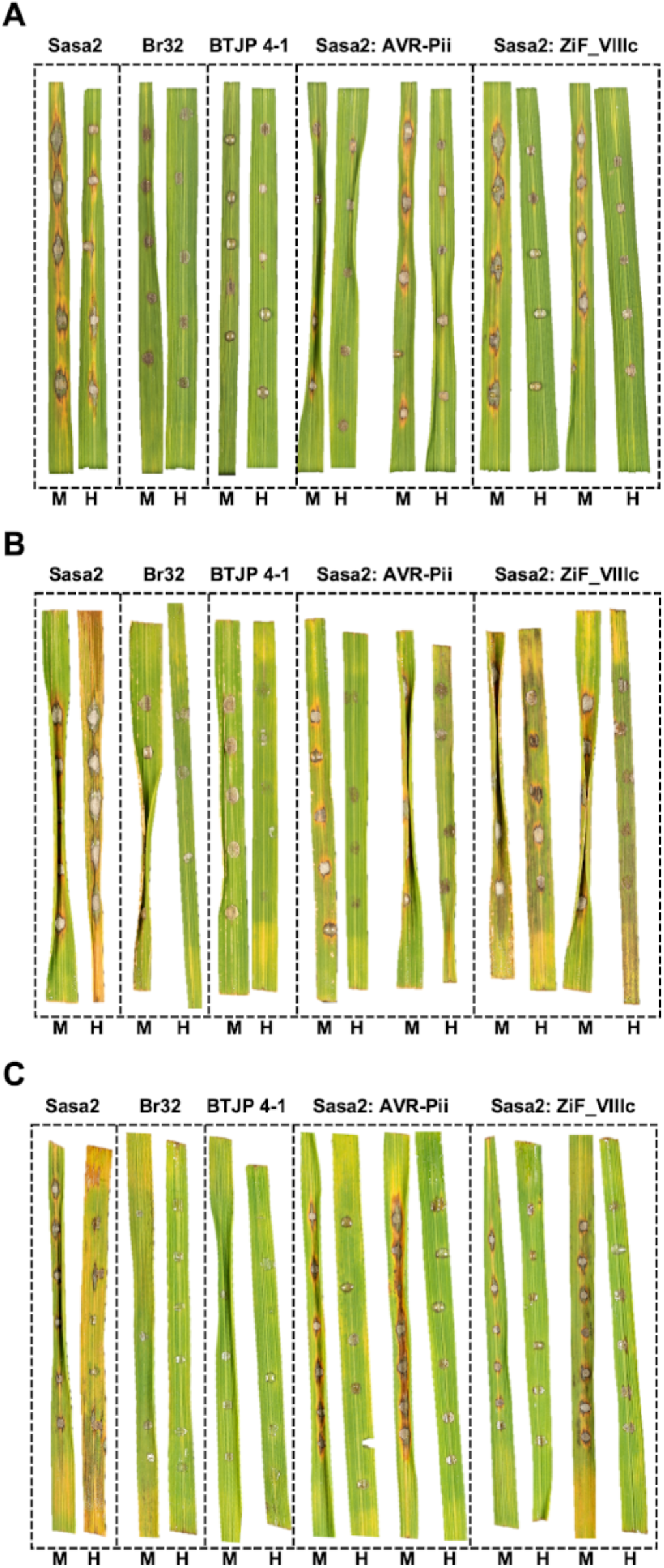
Replicates of the disease resistance assays of Sasa2 harbouring AVR-Pii or ZiF_VIIIc. First **(A),** second **(B)** and third **(C)** replicate of the rice leaf blade spot inoculation assay presented in Figure 4. Transgenic *M. oryzae* Sasa2 harbouring AVR-Pii or ZiF_VIIIc were spotted into rice cultivars Moukoto (Pii-) and Hitomebore (Pii+). The cultivars Moukoto and Hitomebore are denoted by M and H, respectively. Wild-type rice blast isolate Sasa2 and wheat blast Br32 and BTJP 4-1 are included as control.

**Figure S9.**
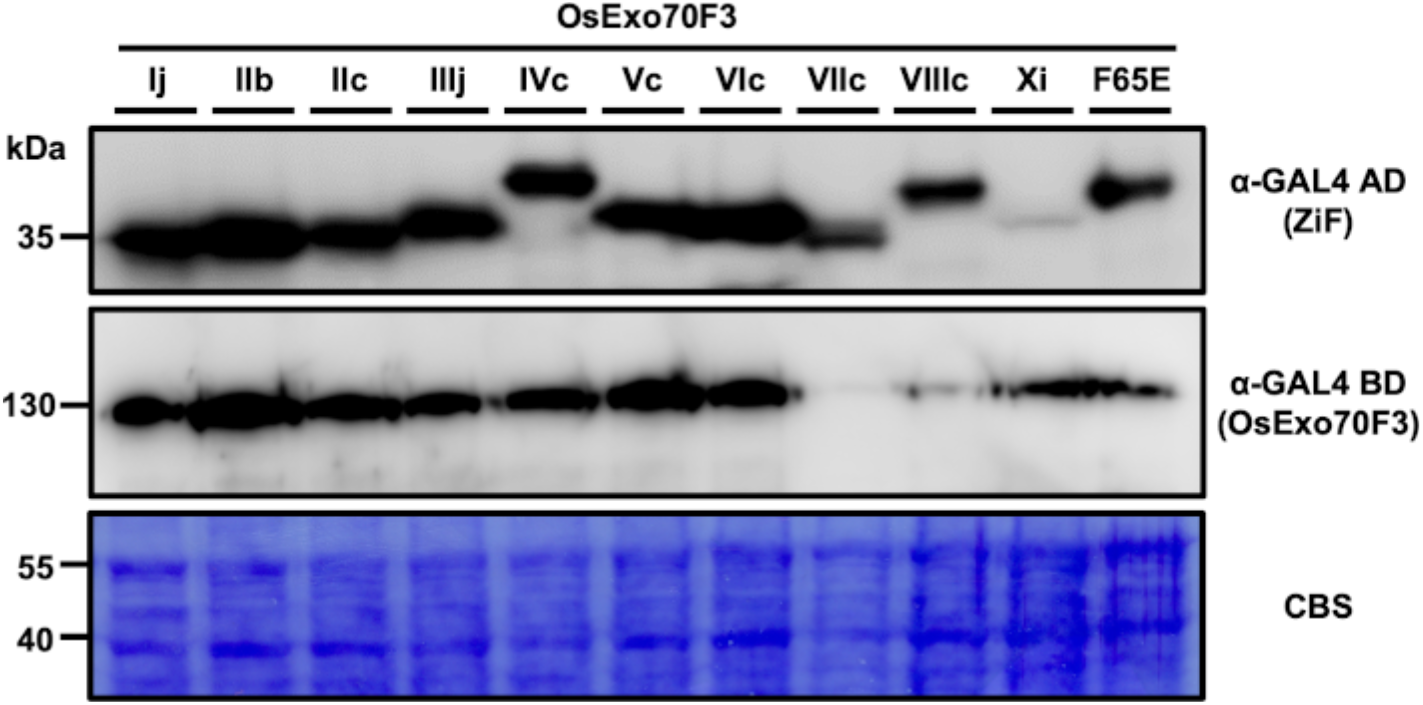
Protein accumulation in rice blast Yeast-Two-Hybrid assay analysed by Western blot. Yeast lysate was probed for the presence of OsExo70F3 using anti-GAL4 binding domain (BD) and the accumulation of selected rice blast ZiF effectors was probed with anti-GAL4 DNA activation domain (AD) antibodies. Total protein extracts were stained with Coomassie Blue Stain (CBS). OsExo70F3 accumulation is consistently lower in positive interactions with ZiF effectors as noticed here and elsewhere in this and previous studies (De la Concepcion et al., 2022).

**Figure S10.**
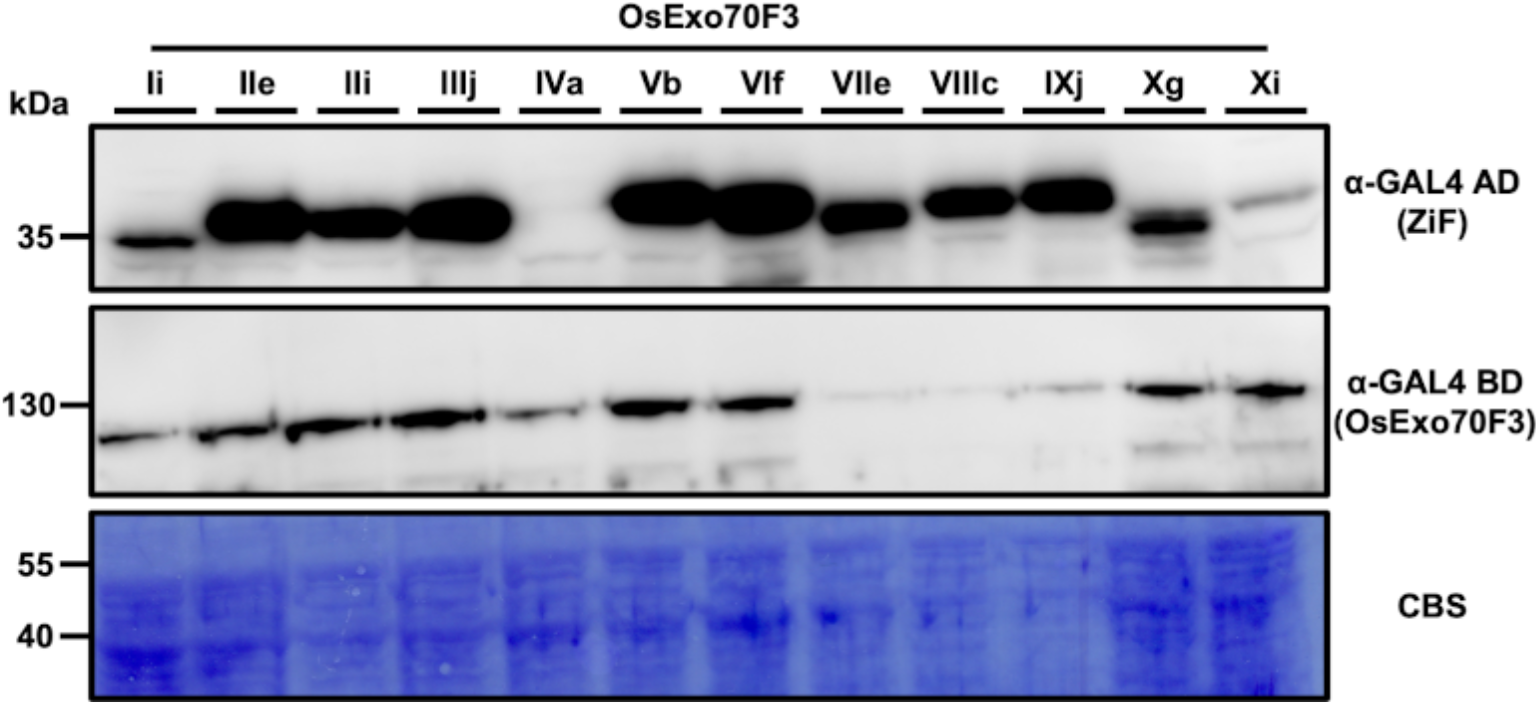
Protein accumulation in wheat blast Yeast-Two-Hybrid assay analysed by Western blot. Yeast lysate was probed for the presence of OsExo70F3 using anti-GAL4 binding domain (BD) while the accumulation of selected ZiF effectors from wheat blast lineages was probed with anti-GAL4 DNA activation domain (AD) antibodies. For technical feasibility, positive and negative controls AVR-Pii and AVR-Pii Phe65Glu were not included in the western blot as their production in yeast cells was tested before (De la Concepcion et al., 2022) and elsewhere in this study. Total protein extracts were stained with Coomassie Blue Stain (CBS). OsExo70F3 accumulation is consistently lower in positive interactions with ZiF effectors as noticed here and elsewhere in this and previous studies (De la Concepcion et al., 2022).

